# Dynamic proteome profiling uncovers age-related impairments in proteostasis and the protective effects of resistance exercise in human skeletal muscle

**DOI:** 10.1101/2025.10.27.684531

**Authors:** Yusuke Nishimura, Krisztina Rudolf, Jennifer Barrett, Richard Kirwan, Kelsie Johnson, Jamie Pugh, Juliette Strauss, José Areta, Connor A. Stead, Daniel Owens, Matthew Jackson, Richard Foster, Sandra Ortega-Martorell, Omid Khaiyat, Claire Stewart, Asangaedem Akpan, Jatin Burniston

## Abstract

A loss of proteostasis is a primary hallmark of ageing that has emerged from mechanistic studies in model organisms, but little is currently known about changes to proteostasis in the muscle of older humans. We used stable isotope labelling (deuterium oxide; D2O) in vivo, and peptide mass spectrometry of muscle samples to investigate differences in proteome dynamics between the muscle of younger (28 ± 5 y; n=4) and older (69 ± 3 y; n=4) men during either habitual activity or resistance exercise training. We quantified the abundance of 1787 proteins and the turnover rate of 1046 proteins in bi-lateral samples of vastus lateralis (n=32 samples total) taken before and after a 15-day program including 5 sessions of unilateral leg-press exercise (3 sets of 10 repetitions at 90% of 10 RM). Our protein abundance profiling revealed a stoichiometric imbalance within the proteostasis network in aged skeletal muscle, including subunits of eIF3, subunits of 40S and 60S ribosomal proteins. The rate of bulk, mixed-protein synthesis was not different between younger and older men, but most ribosomal proteins were less abundant in the muscle of older participants, suggesting ribosomes in older muscle may exhibit increased translational efficiency to maintain similar levels of protein turnover compared to ribosomes in younger muscle. Resistance exercise partially restored age-related disruptions to the proteostasis network. In older skeletal muscle, resistance exercise specifically increased the absolute turnover rate (ATR) of mixed mitochondrial proteins, with increased fractional turnover rate (FTR) of prohibitin 1 (PHB1) and profilin-1 (PROF1), and increased abundance of prohibitin 2 (PHB2). These adaptations may suggest resistance exercise promotes mitochondrial proteostasis by facilitating the synthesis and maintenance of key mitochondrial proteins. Thus, our *Dynamic Proteome Profiling* data provide an impetus for further exploration of the role of proteostasis in maintaining skeletal muscle quality and supports resistance exercise as a potential therapeutic strategy to promote healthy skeletal muscle ageing in humans.

**In Brief:** Nishimura et al. used *Dynamic Proteome Profiling* to uncover whether the distorted proteomic landscape of ageing skeletal muscle is associated with altered turnover of specific proteins. Basal muscle from older men exhibits a divergence in protein abundance between subunits of eIF3 and subunits of 40S and 60S ribosomal proteins, whereas resistance exercise partially restored age-related disruptions in the muscle proteome. In older muscle, protein-specific turnover generally increases after resistance exercise, independent of changes in protein abundance, suggesting improved protein quality and renewal. Created in BioRender. Nishimura, Y. (2025) https://BioRender.com/p2a1aio

**Highlights:**

- *Dynamic Proteome Profiling* in human skeletal muscle ageing
- Ageing alters muscle proteostasis
- Mixed-muscle protein synthesis does not differ between younger and older men
- Resistance exercise increased mitochondrial protein turnover specifically in older muscle
- Protein-specific responses to resistance exercise differed between age groups

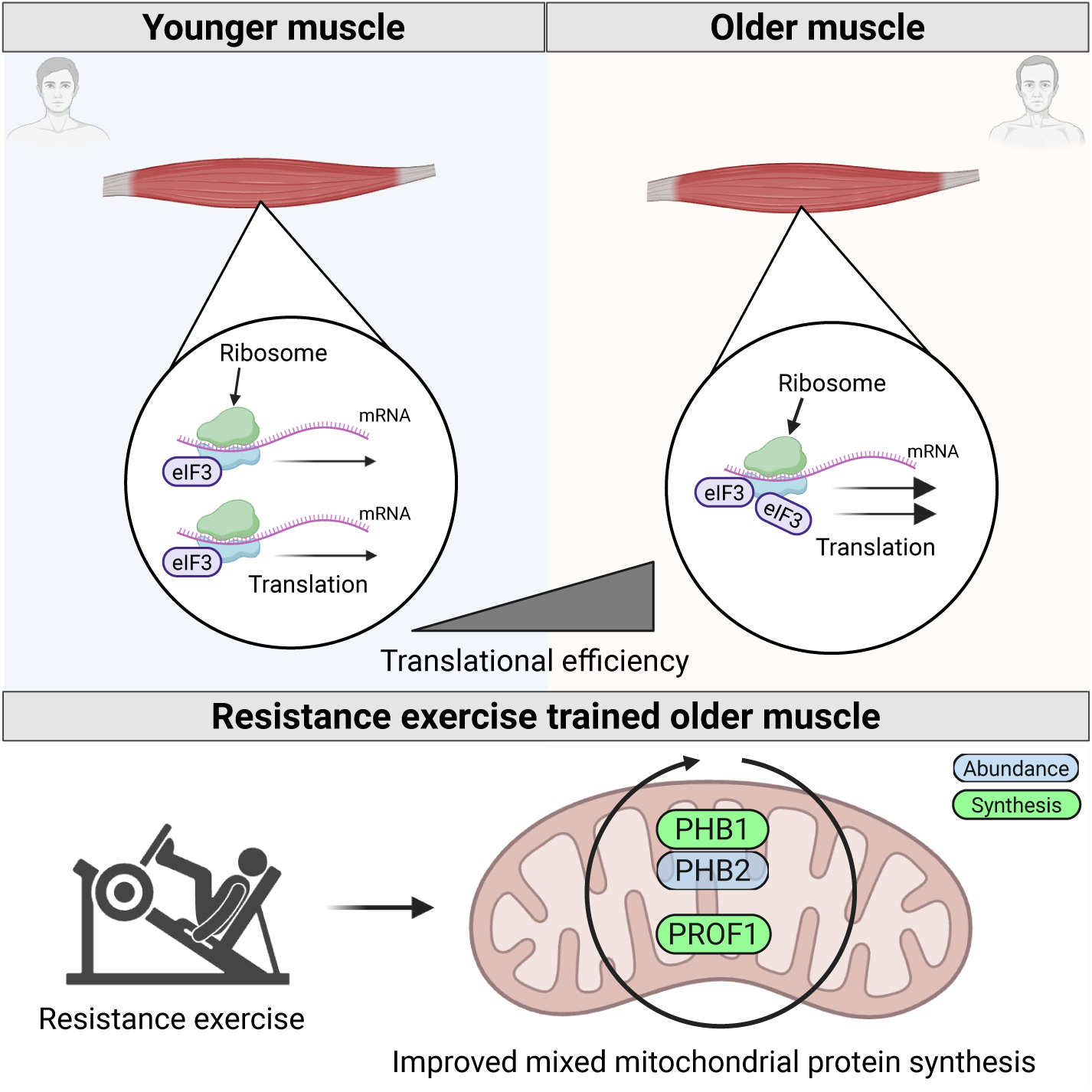

## Introduction

Muscle loss is a hallmark of ageing that is conserved across species, including humans, laboratory rodents and model organisms (1). Older people experience gradual declines in muscle mass and function that negatively impact their risk of chronic disease, resilience to injury, societal/economic contribution and quality-of-life. In older adults, losses in muscle strength occur earlier and are more precipitous than the decline in muscle mass, and predict all-cause mortality independently of age-related muscle loss (2). People over the age of 60 years that rank in the lower tertile of muscle strength have a 50 % higher all-cause mortality rate than those in the higher tertile (3) and sarcopenia directly diminishes quality-of-life (4). Accordingly, in recent years, the focus of epidemiological studies and clinical practice have shifted toward a broader classification of sarcopenia encompassing the extent of losses in both the strength and quantity of skeletal muscle, and the associated losses in physical performance (5). A greater focus on the function and quality of muscle, rather than muscle mass *per se*, may also help align molecular studies of human muscle ageing with mechanistic research in model organisms that highlight losses in proteome quality, i.e. protein homeostasis (proteostasis), as a primary mechanism of ageing (6).

Proteostasis is maintained through protein turnover and the proteostasis network, including the translational machinery, chaperone network, the ubiquitin proteasomal system and autophagic processes (7). The muscle proteome is dominated by relatively large myofibrillar proteins (8) that are exposed to mechanical stress, which creates a high demand for proteostasis machinery to maintain proteome health in skeletal muscle (9). Furthermore, postmitotic tissues such as skeletal muscle are particularly prone to age-related increases in translational error rate (Bottger et al., 2025). The accumulation of erroneously translated, misfolded, or incomplete proteins contribute to losses in proteostasis and present a burden to the proteostasis network (6, 10). Losses in proteostasis are associated with lesser capacity for refolding or proteolysis of damaged proteins, which can result in protein aggregation and subsequent proteotoxicity (11). In model organisms, including *C elegans*, *D melanogaster*, laboratory rodents and primates a loss in proteostasis is a key mechanism of muscle ageing (12). In humans, mitochondrial dysfunction and selective atrophy or loss of fast-contracting myofibres are the currently acknowledged hallmarks of muscle ageing (13) and age-related losses in proteostasis have not yet been comprehensively studied.

Muscle loss is a biomarker of ageing in *C elegans* (14) and older animals exhibit changes to the proteome, including declines in the abundance of mitochondrial and ribosomal proteins, and increases in the abundance of protein chaperones and proteasomal subunits (15). Pulse-chase experiments using stable isotope labelled amino acids (e.g. dynamic SILAC), found the majority (∼70 %) of 546 proteins studied in nematodes decline in turnover rate but 10 % proteins have higher rates of turnover in older animals (16). In particular, the turnover of proteins associated with translation becomes more heterogeneous with age, whereas the rate of turnover of proteins associated with degradative processes is maintained in older animals (16). Studies on human muscle protein dynamics are currently limited to acute measurements on the synthesis rate of mixed-proteins under specific contexts (e.g. fasted or in the immediate period after feeding or exercise). Typically, no difference in mixed-protein synthesis rate is evident between the muscle of younger and older adults under basal conditions (17–19) but older muscle is less sensitive to anabolic stimuli such as feeding (20) or exercise (21), a phenomenon commonly known as ‘anabolic resistance.’ However, when a broad age range (19-87 years) of adults were studied, there was a significant negative correlation between age and muscle protein turnover, and the turnover of mixed-protein in human muscle declined by ∼3.5% per decade (22). Mixed-protein data are a composite of hundreds of different proteins that exhibit a broad range of individual protein turnover rates (23). Therefore, studies on mixed-protein samples overlook the wide heterogeneity of proteins in muscle and age-dependent differences in muscle proteome profile. Moreover, particular proteins exhibit different rates of turnover and respond differently to exercise interventions (24–27).

To date, proteomic analyses on human skeletal muscle ageing are limited to just five studies that compared differences between younger vs older (28–30) adults or investigated the association between age and the proteome across a wide range of the adult lifespan (31, 32). Furthermore, only two proteomic studies incorporated an exercise intervention alongside ageing comparisons (29, 33). Ubaida-Mohien et al. (31) reports approximately 29 % of the muscle proteome (4380 proteins studied) changes with age. The majority (70 %) of proteins that exhibited a significant effect of ageing increased in abundance in older muscles and were associated with genomic maintenance, transcriptional regulators, splicing, neuromuscular junction, proteostasis, senescence and immune function. Proteins that decreased with age were associated with muscle contraction, mitochondrial metabolism and ribosomal function. The notable differences in the abundance profiles of proteins between younger and older skeletal muscle raises challenges to interpreting mixed-protein synthesis data, which cannot distinguish protein-specific changes in turnover rate. We hypothesize, ageing impairs the turnover of specific muscle proteins, but this critical nuance is lost in traditional mixed-protein analyses, in which average turnover rates across hundreds of individual proteins obscure individual responses.

In this study, we used *Dynamic Proteome Profiling* (25) to offer novel insights into human skeletal muscle ageing and the protective effects of resistance exercise. *Dynamic Proteome Profiling* provides protein-by-protein data on both turnover rates and abundance of muscle proteins, providing detailed insights into the two central components of proteostasis: the renewal rate of proteins and the abundance profile of key proteostasis regulators (34). Resistance exercise stimulates skeletal muscle repair and remodeling, counteracting sarcopenia while improving metabolic health and functional capacity (35). In young adults, *Dynamic Proteome Profiling* highlighted previously unappreciated patterns of change, including instances where the abundance of a protein decreases despite and an increase in synthesis rate after resistance exercise (25). In contrast, some proteins may not change in abundance but their rate of turnover is enhanced after exercise (24). In the context of muscle disease (36), some proteins that are more abundant in the disease state also have lower rates of turnover. In the current work, we report that age-related differences in the muscle proteome abundance profile are associated with different dynamic muscle proteome response to resistance exercise between older and younger males, which is critical for optimizing interventions to maintain muscle health with ageing.

## Materials and Methods

### Experimental design

In Fig. 1 A and Fig. 1 B, an overview of the experimental design and protocol is provided, which consisted of a mixed multi-factor design to investigate interactions between age (between subject factor: younger versus older), time (within-subject factor: pre versus post), and training status (within subject factor: exercise versus control leg). Eight healthy male participants consisting of four younger and four older adults were recruited for this study from the Merseyside region, UK. Baseline, anthropological and physiological data were collected from all participants 7 days prior to commencing the 14-day period of D2O consumption. Throughout the experimental period, blood samples were collected (day - 1, 1, 8, 15) and muscle samples were obtained before D2O administration (day -1) and after 15 days of D2O consumption. Participants performed unilateral resistance exercise on days 1, 4, 7, 10 and 13.

**Figure 1.**
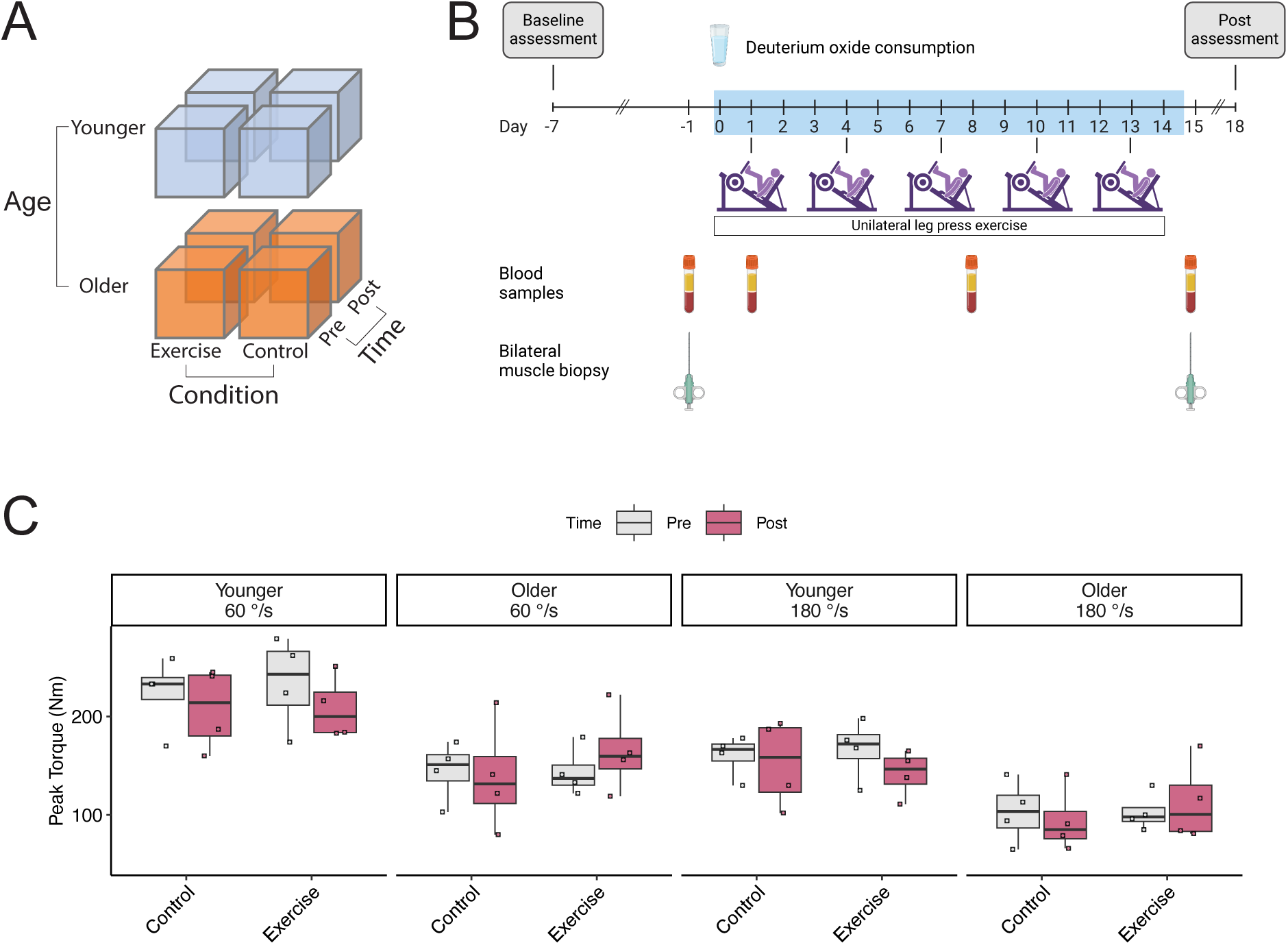
Study design (A) A 3-factor design to investigate interactions between age (younger vs older), time (pre and post), and training status (exercise vs control) using n = 4 subjects/group (n = 32 muscle biopsy samples). (B) Overview of the experimental design. A two-week deuterium oxide (^2^H2O; D2O) labelling experiment was conducted in healthy younger (n = 4) and older men (n = 4). Anthropological and physiological data were obtained at least 7 days before the experiment (day -7, outlined in Table 1). D2O was administered daily (1/20^th^ of their initial dose) over the 14-day labelling period and venous blood samples were collected on days -1, 1, 8, and 15 to calculate body water D2O enrichment. Muscle biopsies of vastus lateralis muscle were conducted on days -1 and 15. All participants performed 5 sessions of unilateral leg press exercise (3 sets of 10 repetitions) at 90% of 10RM interspersed by 3-day recovery periods during the intervention period. (C) Boxplots comparing between limb (Control versus Exercise) and time (Pre versus Post) of knee extensor peak torque (Nm) assessed by an isokinetic dynamometry at two angular velocities.

**Table 1.** Baseline characteristics of participants.

|  | Younger<br>(*n* = 4) | Older<br>(*n* = 4) | P value |
| --- | --- | --- | --- |
| Age (Years) | 28 ± 5 | 69 ± 3 | 7.15e-06*** |
| Height (cm) | 176.4 ± 3 | 173.5 ± 9 | 0.557 |
| Weight (kg) | 76.6 ± 7.5 | 82.3 ± 7.9 | 0.337 |
| BMI (kg/m<sup>2</sup>) | 24.3 ± 2.9 | 27.2 ± 2.6 | 0.12 |
| Fat mass (kg) | 13.0 ± 3.4 | 24.0 ± 6.5 | 0.0236* |
| Fat free mass (kg) | 63.2 ± 5.2 | 57.7 ± 5.7 | 0.201 |
| Percent fat free mass (%) | 83.2 ± 3 | 70.8 ± 6.5 | 0.0147* |
| Visceral Adipose Tissue (l) | 0.53 ± 0.43 | 3.05 ± 0.41 | 0.000152*** |
| Waist Circumference (m) | 0.798 ± 0.046 | 0.93 ± 0.045 | 0.00629** |
| Whole-body skeletal muscle mass (kg) | 31 ± 2.7 | 27.5 ± 3.2 | 0.14 |
| Sit-to-stand (s) | 6.47 ± 1.65 | 7.67 ± 1.92 | 0.379 |
| Knee extension peak torque (60 deg s<sup>-1</sup>, Nm) | 234.75 ± 46.57 | 143.75 ± 24.76 | 0.0136* |
| Knee extension peak torque (180 deg s<sup>-1</sup>, Nm) | 166.75 ± 30.59 | 102.75 ± 19.24 | 0.0122* |
| Systolic blood pressure (mmHg) | 123.5 ± 12 | 147.3 ± 21.2 | 0.0992 |
| Diastolic blood pressure (mmHg) | 66.8 ± 9.2 | 82.5 ± 1.9 | 0.0152* |
| Fasting glucose (mmol/L) | 4.4 ± 0.26 | 5.2 ± 0.26 | 0.00611** |
| Fasting insulin (μU/mL) | 30.7 ± 8.3 | 71.2 ± 11 | 0.000912*** |
| HOMA | 1.025 ± 0.32 | 2.75 ± 0.52 | 0.00132** |

Physical and physiological characteristics measured during pre-experimental testing of healthy younger and older adults. Data presented as mean ± standard deviation. \*P < 0.05; \*\*P < 0.01; \*\*\*P < 0.001

### Participants

Following ethical approval and informed consent, only males that exercised regularly (>3 bouts of endurance-/resistance-based training per week) and were aged between either 25-35 years (younger group) or 65-75 years (older group) were recruited. Exclusion criteria included: previous diagnosis of chronic disease including ischemic heart disease, diabetes mellitus, myopathy or neuromuscular disorder, currently prescribed anticoagulant medications, unwell (e.g. cold or flu) or currently enrolled in any other research study. Volunteers were provided oral and written information regarding the aims, procedures, and potential risks of the study and written informed consent was obtained before participation. The study received approval from the Research Ethics Committee of Liverpool John Moores University (#22/SPS/075) and complied with the Declaration of Helsinki, except for registration in a publicly accessible database.

### Anthropometric and physiological characterization

Anthropometric measurements, including height, weight and abdominal circumference, were recorded using a standard scale and measuring tape. Body composition, comprising fat-free mass, fat mass, and body water content, was assessed using bioelectrical impedance analysis (BIA) (BIA 101 System Analyzer, Akern, Florence, Italy), according to the manufacturer’s guidelines. Functional ability was assessed using the five-times sit-to-stand test, according to standard guidelines (37) and handgrip strength was measured using a digital dynamometer (TKK 5401 Grip-D, Takei, Niigata, Japan) (38), in accordance with the protocol established by Yu et al. (39).

Knee extensor peak torque was assessed using an isokinetic dynamometer (Humac NORM; CSMi, Stoughton, MA, USA), which has been previously validated as a reliable assessment of muscle force output (40). Participants were seated with their hips flexed at 90° and secured with straps to isolate knee extensor activation. Following a standardized warm-up comprising four submaximal repetitions, participants executed four maximal knee extension and flexion movements at two angular velocities: 1.05 rad·s⁻¹ (60°·s⁻¹) and 3.14 rad·s⁻¹ (180°·s⁻¹), with a 3-minute recovery period implemented between trials. The highest recorded peak torque (Nm) during knee extension was analyzed.

### Unilateral leg-press exercise

Participants undertook a supervised training session of unilateral leg-press exercise at a standardized time of day. During each session, participants performed a warm-up consisting of 5 repetitions at 70 % of 10RM. Participants performed three sets of 10 repetitions at a lifting intensity of 90 % of 10 RM with 2- to 3-min rest between sets.

### Muscle sample collection

Muscle samples were obtained from the vastus lateralis of both the dominant and non-dominant legs following an overnight fast (>10 h) on Day -1 and Day 15 of the protocol. Local anaesthesia (1 % Marcaine, ∼2 mL) was administered and muscle samples (∼100 mg) were collected using the conchotome technique (41). Muscle samples were immediately cleaned of blood, excess fat and connective tissue then snap-frozen in liquid nitrogen and stored at -80°C for subsequent analysis.

### Blood sample collection

Venous blood samples (7 mL plasma and 7 mL serum) were drawn from the antecubital fossa following an overnight fast at four-time points throughout the protocol (Fig.1 B). Samples were collected in vacutainers containing sodium fluoride, EDTA, or silica, then centrifuged (1000 × g, 4°C, 10 min). Plasma and sera were aliquoted and stored at -80°C until further analyses.

Fasting plasma glucose levels were determined using an ILab-600 semi-automatic spectrophotometric analyzer with commercial assay kits (Instrumentation Laboratory Ltd, UK). Plasma insulin was measured with a direct insulin ELISA kit (Invitrogen, UK). Insulin sensitivity was evaluated via the Homeostatic Model Assessment (HOMA) and the Matsuda Insulin Sensitivity Index (42).

### Stable isotope labelling protocol and measurement of body water D2O Enrichment

Biosynthetic labelling of newly synthesized proteins was achieved through the oral consumption of deuterium oxide (D₂O) (Sigma-Aldrich, UK). The dosing protocol was personalized based on each participant’s total body water, measured using BIA. On day 0, participants ingested an initial D₂O dose equivalent to 1.5 % of their total body water content distributed across four boluses at approximately two-hour intervals to minimize side effects (e.g., dizziness and nausea). During the subsequent 14-day labelling period, participants consumed a daily maintenance dose equivalent to one-twentieth of the initial dose to account for D₂O loss, given its estimated 10-day half-life in humans.

Body water enrichment of D2O was measured in plasma samples against external standards that were constructed by adding D2O to PBS over the range from 0.0 to 5.0 % in 0.5 % increments. D2O enrichment of aqueous solutions was determined by gas chromatography-mass spectrometry after exchange with acetone, as reported previously (25).

### Preparation of muscle for proteomic analysis

Muscle samples were ground in liquid nitrogen, then homogenized on ice in 10 volumes of 1 % Triton X-100, 50 mM Tris, pH 7.4 (including complete protease inhibitor; Roche Diagnostics, Lewes, United Kingdom) using a PolyTron homogenizer (KINEMATICA, PT 1200 E) followed by sonication (Fisherbrand^TM^). Homogenates were incubated on ice for 15 min, then centrifuged at 1000 x *g*, 4 °C, for 5 min to fractionate myofibrillar (pellet) from soluble (supernatant) proteins. Myofibrillar proteins were resuspended in a half-volume of homogenization buffer followed by centrifuged at 1000 x g, 4 °C, for 5 min. The washed myofibrillar pellet was then solubilized in lysis buffer (7 M urea, 2 M thiourea, 4% CHAPS, 30 mM Tris, pH 8.5). Aliquots of protein were precipitated in 5 volumes of ice-cold acetone, incubated overnight at -20 °C and resuspended in 200 uL of lysis buffer. Proteins were cleared by centrifugation at 12,000 x g, 20 °C, for 20 min. Total protein concentration (μg/μl) was quantified against bovine serum albumin (BSA) standards using the Bradford assay (Thermo Scientific, #23236), according to the manufacturer’s instructions.

Tryptic digestion was performed using the filter-aided sample preparation (FASP) method (43). Aliquots containing 100 µg protein were resuspended in 200 µl of UA buffer (8 M urea, 100 mM Tris, pH 8.5). Proteins were incubated at 37 °C for 15 min in UA buffer containing 100 mM dithiothreitol followed by incubation (20 min at 4 °C) protected from light in UA buffer containing 50 mM iodoacetamide. UA buffer was exchanged for 50 mM ammonium bicarbonate and sequencing-grade trypsin (Promega V5113, Madison, WI, USA) was added at an enzyme to protein ratio of 1:50. Digestion was allowed to proceed at 37 °C overnight, before peptides were collected in 100 μl of 50 mM ammonium bicarbonate containing 0.2 % (v/v) trifluoroacetic acid. Samples containing 4 µg of peptides were de-salted using C18 Zip-tips (Millipore) and eluted in 40% acetonitrile and 0.1% trifluoroacetic acid. Peptide solutions were dried by vacuum centrifugation and peptides were resuspended in 20 µl of loading buffer (2 % (v/v) acetonitrile, 0.1 % (v/v) formic acid) containing 10 fmol/μl yeast ADH1 (MassPrep standard; Waters Corp., Milford, MA) in preparation for LC-MS/MS analysis.

### Liquid Chromatography- tandem mass Spectrometry

Data-dependent analyses of digests from muscle protein fractions were performed using an Ultimate 3000 RSLCTM nano system (Thermo Scientific) coupled to a Q-Exactive orbitrap mass spectrometer (Thermo Scientific). Samples (4.5 μL corresponding to 900 ng of peptides) were loaded on to the trapping column (Thermo Scientific, PepMap100, 5 μm C18, 300 μm X 5 mm), using ulPickUp injection, for 1 minute at a flow rate of 100 μL/min with 2 % (v/v) acetonitrile and 0.1 % (v/v) formic acid. Samples were resolved on a 500 mm analytical column (Easy-Spray C18 75 μm, 2 μm column) using a gradient of 97.5 % A (0.1 % formic acid) 2.5 % B (79.9 % ACN, 20 % water, 0.1 % formic acid) to 50 % A 50 % B over 150 min at a flow rate of 300 nL/min. Data acquisition included MS1 scans at 70,000 resolution at m/z 200 (AGC set to 3^e6^ ions with a maximum fill time of 240 ms) and data-dependent selection of the top- 10 precursors selected from a mass range of m/z 300-1600. MS2 data were acquired after HCD fragmentation (normalized collision energy 30) in the orbitrap analyzer at 17,500 resolution at m/z 200 (AGC target 5e^4^ ion with a maximum fill time of 80 ms) and a 30 s dynamic exclusion window was used to avoid repeated selection of peptides for MS/MS analysis.

### Label-free quantitation of protein abundance

Progenesis Quantitative Informatics for Proteomics (QI-P; Nonlinear Dynamics, Waters Corp., Newcastle, UK, Version 4.2) was used for label-free quantitation consistent with our previous work (25, 36, 44). Prominent ion features were used as vectors to warp each data set to a common reference chromatogram. An analysis window of 10–130 min and 300–1600 m/z was selected. Log-transformed MS data were normalized by inter-sample abundance ratio, and relative protein abundances were calculated using nonconflicting peptides only. Abundance data were then normalized to the 3 most abundant peptides of yeast ADH1 to derive abundance measurements in fmol/ μg protein (45). MS/MS spectra were exported in Mascot generic format and searched against the Swiss-Prot database restricted to Homo-sapiens (20,272 sequences) using a locally implemented Mascot server (v.2.8; www.matrixscience.com). The enzyme specificity was trypsin with 2 allowed missed cleavages, carbamidomethylation of cysteine (fixed modification) and oxidation of methionine (variable modification). M/Z error tolerances of 10 ppm for peptide ions and 20 ppm for fragment ion spectra were used. Peptide results were filtered to 1% FDR based on decoy search and at least one unique peptide was required to identify each protein. The Mascot output (xml format), restricted to non-homologous protein identifications was recombined with MS profile data in Progenesis.

### Calculation of protein synthesis rates

Mass isotopomer abundance data were extracted from MS spectra using Progenesis Quantitative Informatics (Non-Linear Dynamics, Newcastle, UK). Consistent with previous work (25, 36), the abundances of peptide mass isotopomers were collected over the entire chromatographic peak for each proteotypic peptide that was used for label-free quantitation of protein abundances. Mass isotopomer information was processed using in-house scripts written in Python (version 3.12.4). The incorporation of deuterium into newly synthesized protein was assessed by measuring the increase in the relative isotopomer abundance (RIA) of the m1 mass isotopomer relative to the sum of the m0 and m1 mass isotopomers (Equation 1) that exhibits rise-to-plateau kinetics of an exponential regression (46) as a consequence of biosynthetic labelling of proteins *in vivo*.

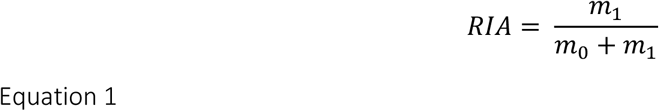

The plateau in RIA (*RIAplateau*) of each peptide was derived (Equation 2) from the total number (*N*) of ^2^H exchangeable H—C bonds in each peptide, which was referenced from standard tables (47), and the difference in the D:H ratio (^2^H/^1^H) between the natural environment (*DHnat*) and the experimental environment (*DHexp*) based on the molar percent enrichment of deuterium in the precursor pool, according to (48).

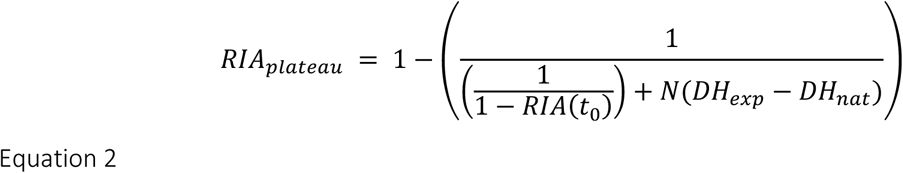

The rate constant of protein degradation (*kdeg)* was calculated (Equation 3) between the beginning (t0) and end (t1) of labelling period. Calculations for exponential regression (rise-to-plateau) kinetics reported in (48) were used and *Kdeg* data were adjusted for differences in protein abundance (P) between the beginning (t0) and end (t1) of labelling period.

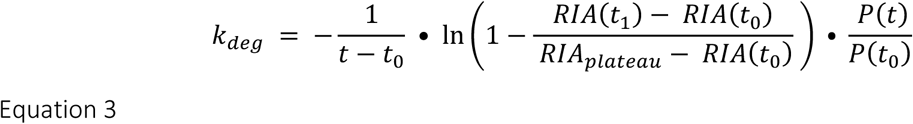

Protein abundance (P; ng/ μg protein) of each protein was calculated from abundance values (fmol/ μg total protein) measured by LFQ multiplied by the molecular weight (MW; kDa) of the protein (referenced from the UniProt Knowledgebase). Absolute turnover rates (ATR; ng/ μg protein/ d) were derived (Equation 4) by multiplying peptide *Kdeg* by the abundance (e.g. ng/ μg protein) of the protein at the end of the labelling period P(t).

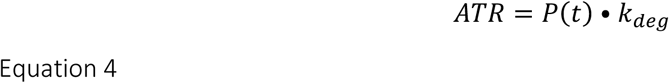

Fractional turnover rates (FTR; %/d) were calculated (Equation 5) from the ratio between ATR (ng/ μg total protein/ d) and the abundance (ng/ μg total protein) of the protein at the end of the labelling period P(t).

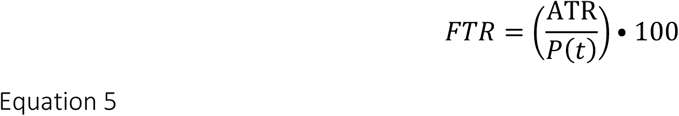

### Statistical and bioinformatic analysis

The experiment was designed to investigate differences in the abundance and turnover rate of proteins between younger and older adults and the response to resistance exercise training. All statistical analysis was performed in R (Version 4.5.1).

Baseline participant characteristics and physiological data were analyzed using between-subjects one-way ANOVA. Proteomic data were analysed by linear mixed-effects models using lme4 (49) to investigate interactions between age (younger versus older) and exercise status (control versus exercised leg) accounting for within-subject measurements at baseline and after the experimental intervention. Accordingly, age and intervention were included as fixed effects and subjects were specified as random effects. Prior to statistical analyses, protein abundance and turnover rate data were filtered to exclude proteins that were not measured in at least N=3 subjects per group or condition.

Baseline differences in protein abundance between younger and older adults were investigated using data averaged across samples from the dominant and non-dominant legs within each participant. Age-related differences in protein turnover rates were investigated using data from the control leg only. Differences in the abundance or turnover rate of proteins between younger and older age groups, at baseline or after the experimental intervention in control and exercised legs were reported as log2 transformed data and statistical significance was set at P < 0.05. Due to the limited number of replicates (N ≥3, per group), a false discovery rate threshold was not set, instead, q values (50) corresponding to proteins a the level of P ≤ 0.05 were reported. In addition to comparisons of protein specific data, the average ATR across all quantified proteins calculated for each participant within in silico-defined myofibrillar and mitochondrial protein sets, providing a mixed-protein ATR estimate for each compartment.

Gene ontology (GO) analyses were performed via Overrepresentation Enrichment Analysis (51) using Gene Ontology enRIchment anaLysis and visuaLizAtion tool (Gorilla) (52). Enrichment of GO terms was considered significant if the Benjamin Hochberg adjusted p value was 0.01, using a study-specific background comprised of all proteins quantified and included in statistical analysis. The coverage of mitochondrial proteins was survey against the Human MitoCarta 3.0 database (53).

## RESULTS

### Participant characteristics

All participants were healthy, had normal anthropological and physiological characteristics (Table 1), and reported partaking in regular exercise and structured physical activity. Body weight and BMI did not differ between the age groups, but older adults had 85 % greater (P = 0.0236) fat mass, and 16 % lesser (P = 0.201) fat-free mass compared to younger adults. Under fasted conditions, older adults had significantly higher blood levels of glucose and insulin (Table 1), indicating impaired whole body glucose metabolism. The HOMA-IR index of older adults (2.75 ± 0.52) was significantly (P = 0.00132) greater than younger adults (1.025 ± 0.32) and was outside the normative range (0.5 – 1.4) (54), which is consistent with expected age-related increases in insulin resistance (55). After consumption of deuterium oxide, deuterium enrichment of the body water compartment rose to a maximum of 1.39 ± 0.05 % in younger and 1.4 ± 0.06 % in older individuals by Day 1 and was maintained above 1% throughout the 15-day experimental period.

Knee extensor peak torque of the dominant and non-dominant leg was significantly greater in younger adults during extension at 60°/s and 180°/s (Fig.1 C), consistent with previous findings (56) in humans. The difference in knee extensor peak torque at 60 °/s between younger adults (234.75 ± 46.57 Nm) and older adults (143.75 ± 24.76 Nm) equates to age-related rate of decline of ∼2.2 Nm/ year, which is in line with expected values (57). The decline in knee extensor strength has critical implications for mobility and independence, particularly among older adults because deficits in strength increase the risk of falls and functional impairments (58).

### Dynamic proteome profiling of human skeletal muscle

Proteomic analyses were conducted on 32 samples of vastus lateralis, encompassing both left and right legs taken prior to (Day -1) and after (Day 15) the experimental intervention in younger (N=4) and older (N=4) men (Fig.1 A, B). In total, 1835 proteins were confidently identified (>1 unique peptide at a false identification threshold of 1 %) and, after filtering to exclude proteins that had >1 missing value per group, the abundances of 1787 proteins were quantified. The median protein sequence coverage was 13.6 % and 92.7 % of peptides used for LFQ had no missed cleavages. The coefficient of variation in protein abundances between the dominant and non-dominant leg of each participant at baseline had a median value of 17.4 % and 7.8 – 35.1 % inter-quartile range, indicating a good level of within-subject reproducibility from independently processed samples.

Further filters were applied to select peptides with clearly resolved envelopes of m0 and m1 mass isotopomers for the calculation of protein FTR. A two-step filtering approach was applied to the FTR data, to ensure data quality and biological relevance. First, proteins were retained only if they had paired measurements (Days -1 and 15) from both the normal-activity control and resistance-exercised leg in each subject, ensuring that comparisons between interventions were based on complete within-subject data. Second, proteins were required to have valid measurements from N ≥ 3 participants within each age group (younger and older). After filtering, FTR data for a total of 1046 proteins were included in the statistical analyses.

### Differences in protein turnover rates and abundance profile between the muscle of younger and older adults at baseline

In the control leg of younger adults, the average (mean ± SD) of protein-specific FTR values was 4.23 ± 2.75 %/d and ranged between 0.08 %/d (laminin subunit alpha-5; LAMA5) and 15.7 %/d (vascular protein sorting 35; VPS35). Myosin heavy chain isoforms were highly abundant in human muscle and the profile of adult muscle isoforms (MYH1, MYH2, and MYH7) did not differ (Two-way ANOVA P > 0.05) between younger and older adults at baseline (Supplemental Fig.1). Most of the highly-abundant muscle proteins had FTR values below the mean value and when protein FTR data were weighted by protein abundance (Fig.2 A) the average turnover rate of protein in young adult muscle was 1.69 %/d. The muscle of younger and older adults may exhibit differences in either or both protein synthesis rate and abundance profile. Therefore, ATR data (ng/ μg protein/ d) combining the abundance and turnover rate of each protein were used to compare mixed-protein averages. Mean ATR (ng/ μg protein/ d) in the control leg was not different (*NS*) between younger (14.65 ± 5.35) and older (15.42 ± 2.77) adults and there were no differences in the distribution profile of protein-specific ATR (Fig.2 B) between younger and of older adults.

**Figure 2.**
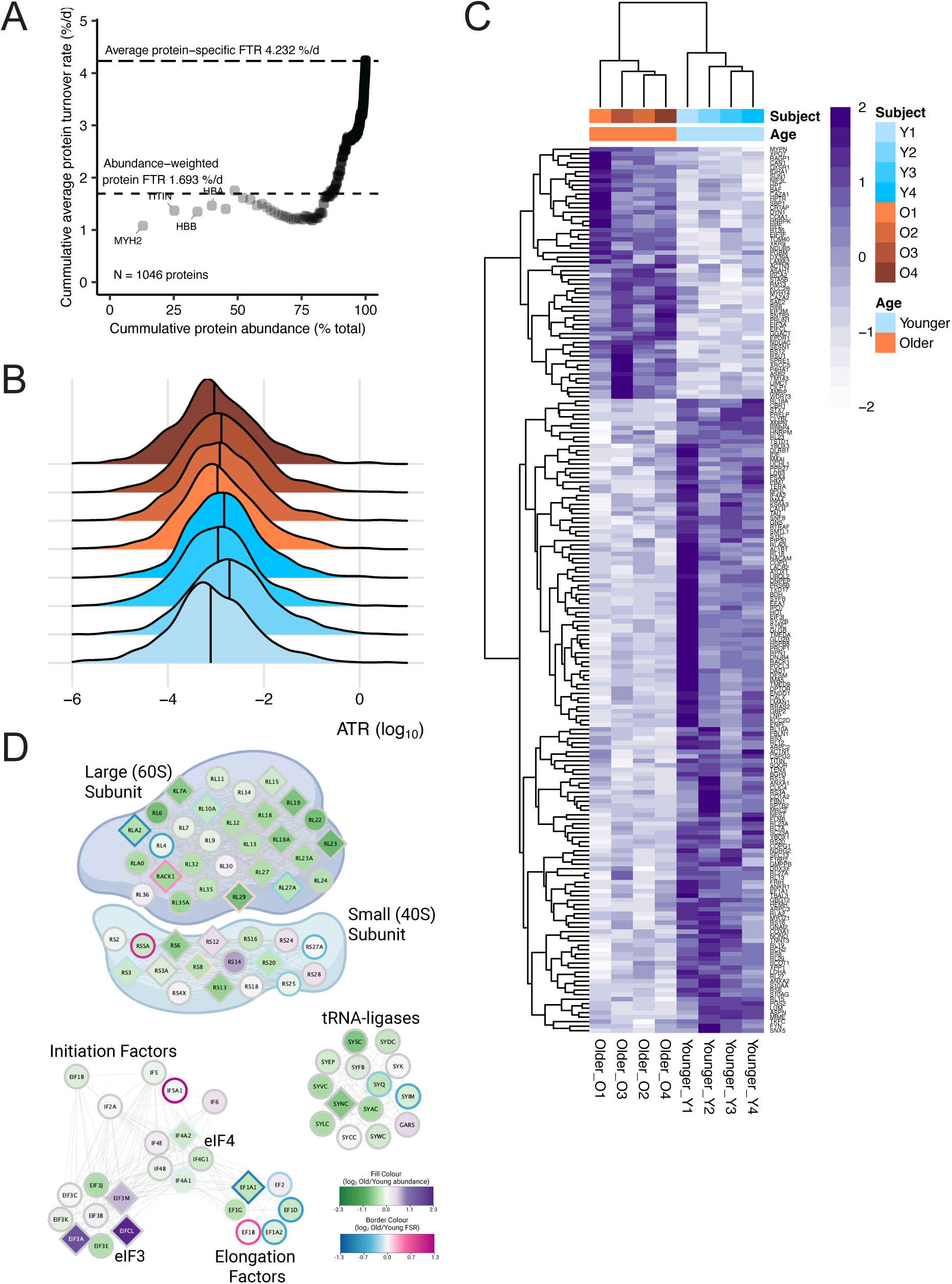
Protein abundance data uncovered a key imbalance in proteostasis network in older skeletal muscle at baseline (A) Cumulative distribution of protein abundance and turnover rate in the control leg of younger men. The x-axis represents the cumulative protein abundance as a percentage of total protein abundance (% total). The y-axis represents the cumulative protein turnover rate (%/d). (B) Density plot of log10 transformed ATR data in the control limb. The vertical black line in each density plot indicates the median ATR for each sample. (C) Heatmap of proteins with significant differences in abundance between younger and older muscle (197 proteins, P < 0.05). Protein abundances were normalized by row and represented by colors according to the scale in the key. Both rows (proteins) and columns (samples) were clustered using hierarchical clustering. (D) Nodes represent proteins organized to their principal protein translation pathway, annotated by their UniProt knowledgebase identifier. Node fill color represents Log2 fold-difference in abundance (Old/ Young) Diamond shaped nodes represent proteins exhibiting significant (P < 0.05) differences in abundance between the age groups at baseline. Node boarder color represents group mean log2 fold-difference in fractional turnover rate (FTR; Old/Young) at baseline. Grey borders indicate missing FTR data.

Protein-specific, differences between the muscle of younger and older adults were investigated by separate analyses on FTR and abundance data, which is necessary to distinguish differences in either or both turnover rate and protein abundance. Twenty-four proteins exhibited differences (P<0.05, q = 0.997) in FTR between the control muscle of younger and older adults. Proteins with particularly high FTR in older muscle, included the mitochondrially encoded ATP synthase F(0) complex subunit alpha (ATP6), apolipoprotein B-100 (APOB) and eukaryotic translation initiation factor 5A-1 (IF5A1). Whereas, myosin-8 (MYH8), protein arginine N-methyltransferase 1 (ANM1) and 4’-phosphopantetheine phosphatase (PANK4) were amongst the proteins with the lowest FTR values in older muscle.

Age-related differences in protein abundance were compared using samples from both left and right legs prior to exercise and revealed 197 significant (P < 0.05) differences, including 63 differences by age at q < 0.15 (Fig. 2 C, Supplemental Table S1). Proteins associated with cytoplasmic translation (GO #0002181), including 11 subunits of the 40S ribosome and 28 subunits of 60S ribosome (Fig.2 D), were significantly enriched amongst the 121 proteins that were less abundant in older compared to younger muscle (main effect of age). The current dataset encompassed 348 of the 1137 mitochondrial proteins annotated in the Human MitoCarta3.0 database (53), including 82%, 100%, 90%, 62%, and 57% of subunits belonging to respiratory chain complexes I-IV, respectively. In accordance with reported (31) age-related declines in human muscle mitochondrial protein abundance, 66 out of 74 subunits of the mitochondrial respiratory complexes quantified were less abundant in older compared to younger muscle.

No gene ontology terms were statistically enriched amongst the 51 proteins that were more abundant in older muscle, but proteins that were significantly more abundant in older muscle included 4 subunits (A, CL, F and M) of eukaryotic Initiation factor 3, small ribosome subunit proteins S12 (RS12), mitochondrial ribosomal subunits RM13 and Heterogeneous nuclear ribonucleoprotein K (HNRPK) which were contrary to the general pattern of lesser abundance of proteins associated with ribosomal translation in older muscle. Basement membrane (Laminin subunit alpha-2, collagen alpha-1 (XVIII) PGBM and cytoskeletal proteins, myopalladin, actinin-3 including actin capping proteins, calpain and transmembrane protein 143 were amongst the proteins more abundant in older muscle. Proteins linked to muscle mass, including Ankyrin repeat and SOCS box protein 2 (ASB2) and Sestrin-1 (SESN1) were more abundant in older muscle. SESN1 is a known negator of mTORC1 (59). Overexpression of ASB2β induces muscle atrophy and a decline in force production, whereas knockdown of ASB2β slightly induces muscle hypertrophy (60).

### Differences in the muscle response to resistance exercise in younger and older adults

Compared to the control leg, mixed-protein ATR tended (*NS*) to be greater in the resistance trained leg, which performed 5 sessions of leg press exercise during the 15-day investigation period (Fig. 3 A, B, C). Compartment-specific responses were investigated by assigning proteins to mitochondrial (231 proteins, Fig. 3 B) and myofibrillar (202 proteins Fig. 3 C) components *in silico* based on gene ontology annotations. Although not statistically significant, myofibrillar proteins tended to exhibit a greater synthetic response to exercise in both younger and older adults (Fig. 3 C), whereas mitochondrial proteins tended to exhibit a response to resistance exercise only in older adults (Fig. 3 B). Consistent with our earlier use of *Dynamic Proteome Profiling* (25), the FTR of the fast-twitch IIa isoform of myosin heavy chain (MYH2) was specifically increased by resistance exercise in young adults and, herein, we found this response to exercise is conserved in older adults (significant main effect of exercise, Fig.3 D). In addition, resistance exercise increased synthesis of the peri-natal isoform, MYH8, in both younger and older muscles (Fig.3 D). MYH8 is a neonatal myosin heavy chain associated with muscle development and regeneration (61). Exercise increased MYH8 protein abundance in younger muscle, whereas in older muscle, the abundance of MYH8 decreased (significant interaction effect, Fig.3 E), which may indicate inefficient muscle remodeling in older muscles.

**Figure 3.**
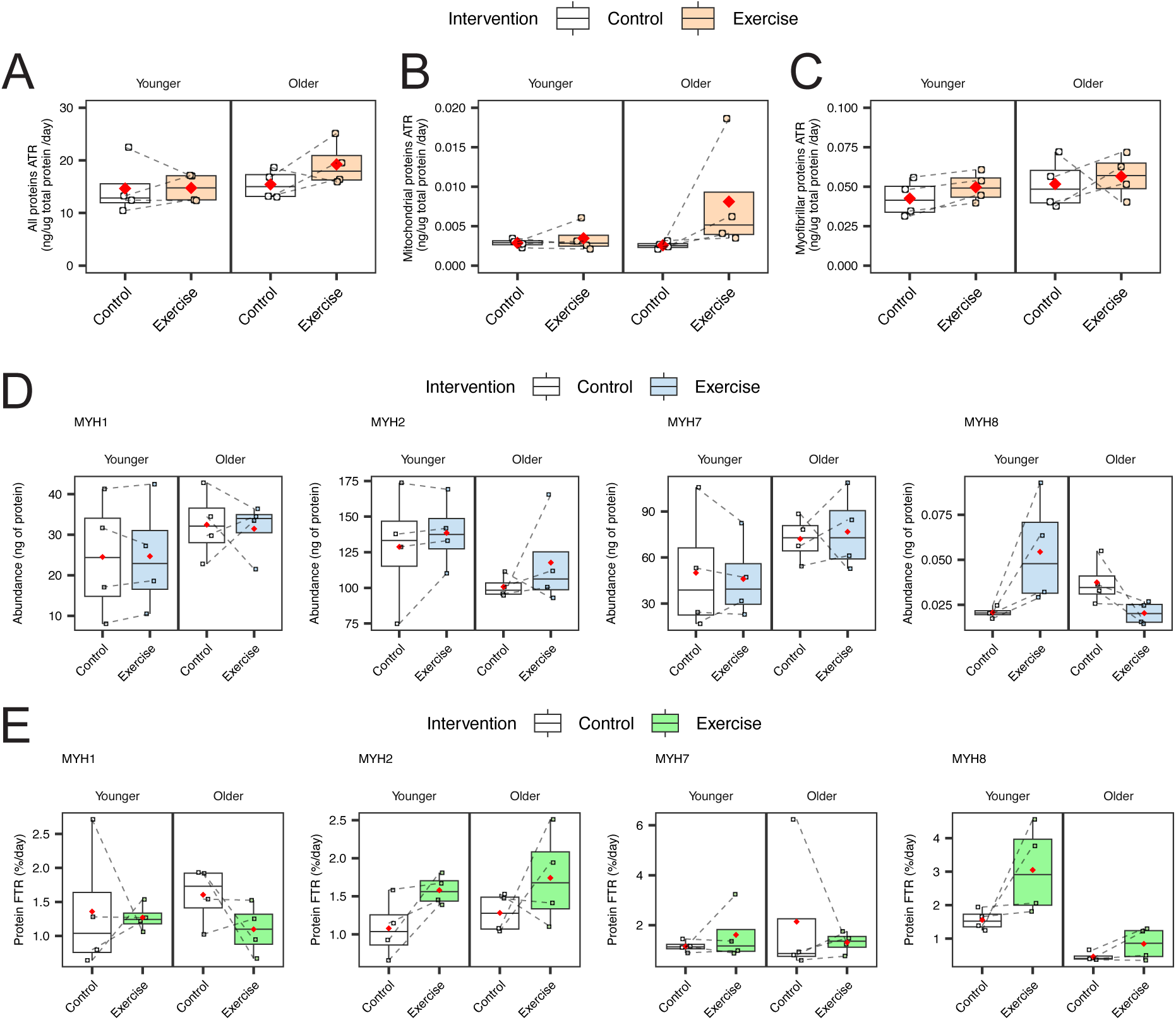
Compartment-specific ATR and myosin heavy chain isoform-specific responses to resistance exercise Absolute turnover rate (ATR) of proteins annotated into (A) total (1045 proteins), (B) mitochondrial (231 proteins), and (C) myofibrillar (202 proteins) *in silico*. The box plot represents the interquartile range (IQR; 25th–75th percentile), with the horizontal line indicating the median. Whiskers extend to the minimum and maximum values within 1.5× IQR. ATR (ng/μg of protein/day) values for individual subjects are shown as individual data points. Dashed lines connect values from the same subject between intervention (Control and Exercise). The red diamond represents the mean. Boxplot of myosin heavy chain isoform-specific proteins comparing age (Younger vs Older) and intervention (Control vs Exercise) in FTR (D) and abundance (E).

Non-targeted inspection of FTR data highlighted distinct patterns (Fig. 4, Supplemental Table S2) amongst the changes in protein-specific responses to resistance training. Fifty-one proteins exhibited an interaction (Fig. 4 A, C, D, P < 0.05, q = 0.87, pink data points) between age (younger vs older adult) and intervention (control vs resistance exercise). Twenty-seven of these proteins were localized in the top left quadrant, where FTR was decreased by exercise in younger muscle but increased in older muscle (Fig. 4 C). Eight proteins (COX6C, PHB, M2OM, UCRI, COQ7, AOFA, MCAT, TIM9) that occupied the top left quadrant were annotated in the Human MitoCarta database, which is consistent with the trend in ATR exhibited by mixed mitochondrial proteins (Fig. 3 B). Several cytoskeletal and structural proteins were also localized in the top left quadrant, including CAZA2, TAGL2, PROF1, SPEG, SNTA1, MFAP5, CORO6, and CAD13. The greater FTR of cytoskeletal proteins during resistance exercise specifically in the muscle of older adults may reflect a greater requirement for structural repair or adaptation. Accordingly, proteins involved in the ubiquitin proteasome system (RD23A, UBP14, UBC12, PHP14) were similarly localized in the top left quadrant.

**Figure 4.**
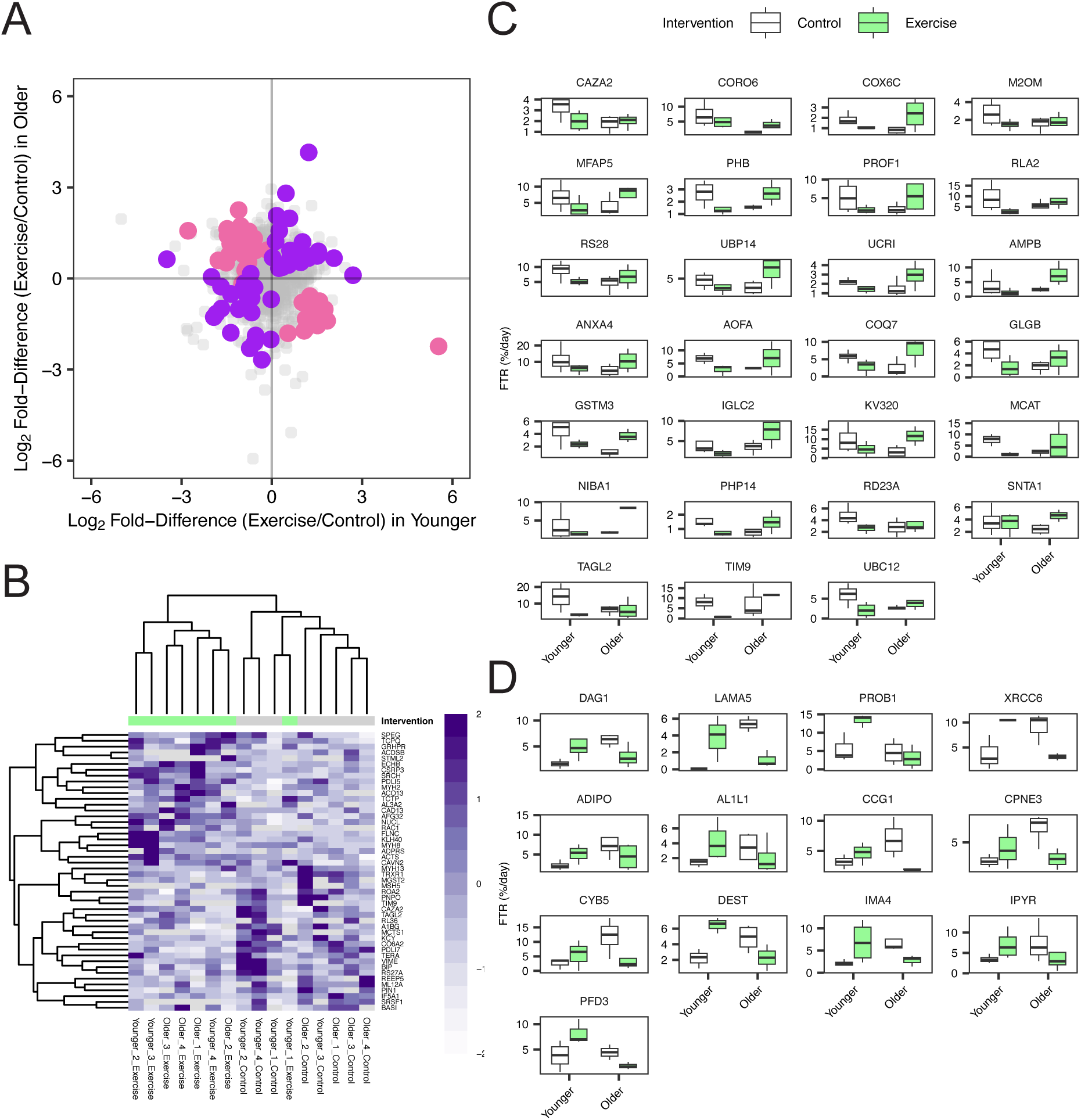
Changes of protein-specific FTR in response to resistance exercise (A) Scatter plot comparing the Log2 Fold-Difference (Exercise/Control) between FTR of Younger (x-axis) and Older (y-axis) adults. Proteins that exhibited an interaction effect (P<0.05) are highlighted in pink and proteins that exhibit a main effect of intervention (P<0.05) are highlighted in purple. (B) Heatmap of proteins with a significant main effect of intervention (48 proteins, P < 0.05). Protein-specific FTR was normalized by row and represented by colors according to the scale in the key. Box plots of 27 proteins with an interaction effect localized in the top left quadrant (C) and 13 proteins with an interaction effect localized in the bottom right quadrant (D).

Thirteen proteins associated with remodeling of the muscle extracellular matrix (ECM) and muscle cytoskeleton were localized in the bottom right quadrant, which indicates the FTR of these proteins was increased by exercise in younger but decreased in older muscle, Fig.4 D). The ECM and cytoskeletal-related proteins included laminin subunit alpha-5 (LAMA5), destrin (DEST) and dystroglycan 1 (DAG1). In addition, prefoldin subunit 3 (PFD3) and the transitional endoplasmic reticulum ATPase (TERA) were also localized in the bottom right quadrant. PFD3 transfers unfolded actin to cytosolic chaperonin and is necessary for the correct folding of actin and tubulin (62). TERA is a AAA+ ATPase involved in endoplasmic reticulum-associated protein degradation (ERAD) and removes misfolded or damaged proteins from the endoplasmic reticulum (63).

Protein abundances also exhibited interaction effects between age (younger versus older adult) and time (baseline versus post-intervention) in the exercised leg (Fig. 5, Supplemental Table S3). Eight proteins, including the mitochondrial inner membrane components, TIM13 and PHB2, decreased in younger and increased in older muscle in response to resistance exercise (Fig. 5 B). In contrast, proteins that had been more abundant in older than younger muscle at baseline, including ASB2, EIFCL, MYPN, NDUAC, NDUB5, and TOM40, decreased in older muscle in response to resistance exercise training (Fig. 5 C). In addition, key muscle proteins such as perinatal myosin heavy chain (MYH8) and lamtor 1 (LTOR1), which tethers the Ragulator complex to the lysosomal membrane, were also found in bottom right quadrant (Fig. 5 C).

**Figure 5.**
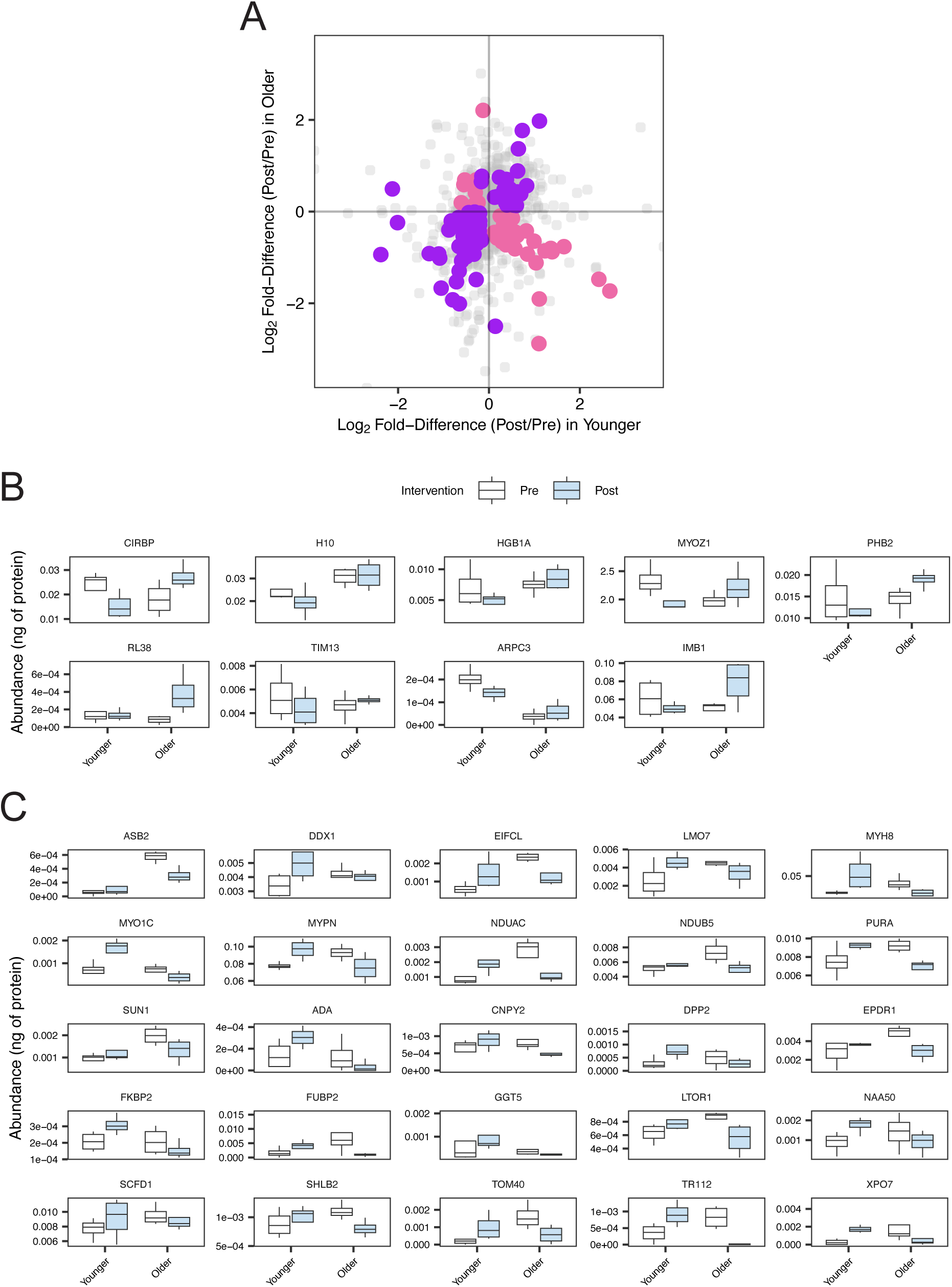
Changes of protein abundance in response to resistance exercise (A) Scatter plot comparing the differences in the Log2 Fold-Difference (Post/Pre in resistance exercise limb) between protein abundance of Younger (x-axis) and Older (y-axis) muscles. Proteins that exhibited an interaction effect (P<0.05) are highlighted in pink and proteins that exhibit a main effect of intervention (P<0.05) are highlighted in purple. Box plots of 9 proteins with an interaction effect localized in the top left quadrant (B) and 25 proteins with an interaction effect localized in the bottom right quadrant (C). Box plots of proteins exhibited a main effect of intervention is presented in Supplemental Fig. 2.

### Similarities in the muscle response to resistance exercise in younger and older adults

Forty-eight proteins exhibited FTR responses to resistance exercise that were similar in younger and older adults (P<0.05, main effect of intervention). Twenty-three of those proteins increased synthesis (Fig. 4 B) and included core myofibrillar proteins (MYH2, MYH8, Fig. 3 D), sarcomeric and Z-disc associated proteins such as FLNC, SPEG, CSRP3, CAPZA2, and KLHL40 and cytoskeletal organization proteins such as PDLI5, CAD13, CAVN2, GRHPR, and AL3A2 also showed elevated synthesis. Mitochondrial proteins including AFG32, ACDSB, ACO13, ECHB, and STML2 also had higher FTR values in exercised muscle and RAC1, which regulates insulin-dependent glucose uptake and contraction-stimulated GLUT4 translocation (64) increased in synthesis after resistance exercise in both younger and older adults. Conversely, the turnover rates of endoplasmic reticulum chaperone BiP (BIP), Myosin regulatory light chain 12A (ML12A), collagen alpha-2 (VI) chain (CO6A2), eukaryotic translation initiation factor 5A-1 (IF5A1), myosin 13 (MYH13), PDZ and LIM domain protein 7(PDLI7), ubiquitin-40S ribosomal protein S27a (RS27A), Serine/arginine-rich splicing factor 1 (SRSF1), and vimentin (VIME) decreased in response to resistance exercise training (Fig. 4 B). Some exercise-induced changes in protein abundance were also common amongst younger and older adults. Subunits of the mitochondrial electron transport chain and proteasome (PSMD7, PSB3, PSB4, PRS6B, PRS8) decreased after resistance exercise, whereas protein subunits of the ribosome, including RL14 and RS12 and cytoskeletal proteins, including FLNB, SPTAN1, VIM were increased by resistance exercise (Supplemental Fig.2).

### Divergent dynamic proteome responses to resistance exercise in younger versus older adults

The separate analyses of age and intervention differences in protein FTR (Fig. 4) and protein abundances (Fig. 5) were combined to illustrate differences in the pattern of the muscle dynamic proteome response to exercise between younger (Fig. 6 A) and older (Fig. 6 B) men. The proteins categorized into four quadrants in younger muscle (Fig. 6 A) were mapped based on their log2 Fold- Difference (Exercise/Control) in abundance and FTR in older muscle (Fig. 6 B) to illustrate age-related differences in resistance exercise response. Muscle responses to resistance exercise in young adults included proteins that decreased in abundance and increased in FTR (Fig. 6 A; Q1, top-left), increased in abundance and increased in FTR (Fig. 6 A; Q2, top-right), decreased in abundance and decreased in FTR (Fig. 6 A; Q3, bottom-left) and increased in abundance and decreased in FTR (Fig. 6 A; Q4, bottom-right). Proteins that occupied Q4 (increased in abundance but decreased in FTR) in response to exercise in younger adults were redistributed toward Q2 (increased in both abundance and FTR) in older muscle (Fig. 6 B). In older muscle, proteins that occupied Q2 were characterized by actin binding (GO:0003779) and actinin binding (GO:0042805). Proteins that occupied Q3 (decreased in both abundance and FTR) in response to exercise in younger adults were redistributed toward Q1 (decreased in abundance and increased in FTR) in older muscle (Fig. 6 B) and these proteins were characterized by mitochondrion (GO:0005739), tricarboxylic acid cycle (GO:0006099), electron transport chain (GO:0022900).

**Figure 6.**
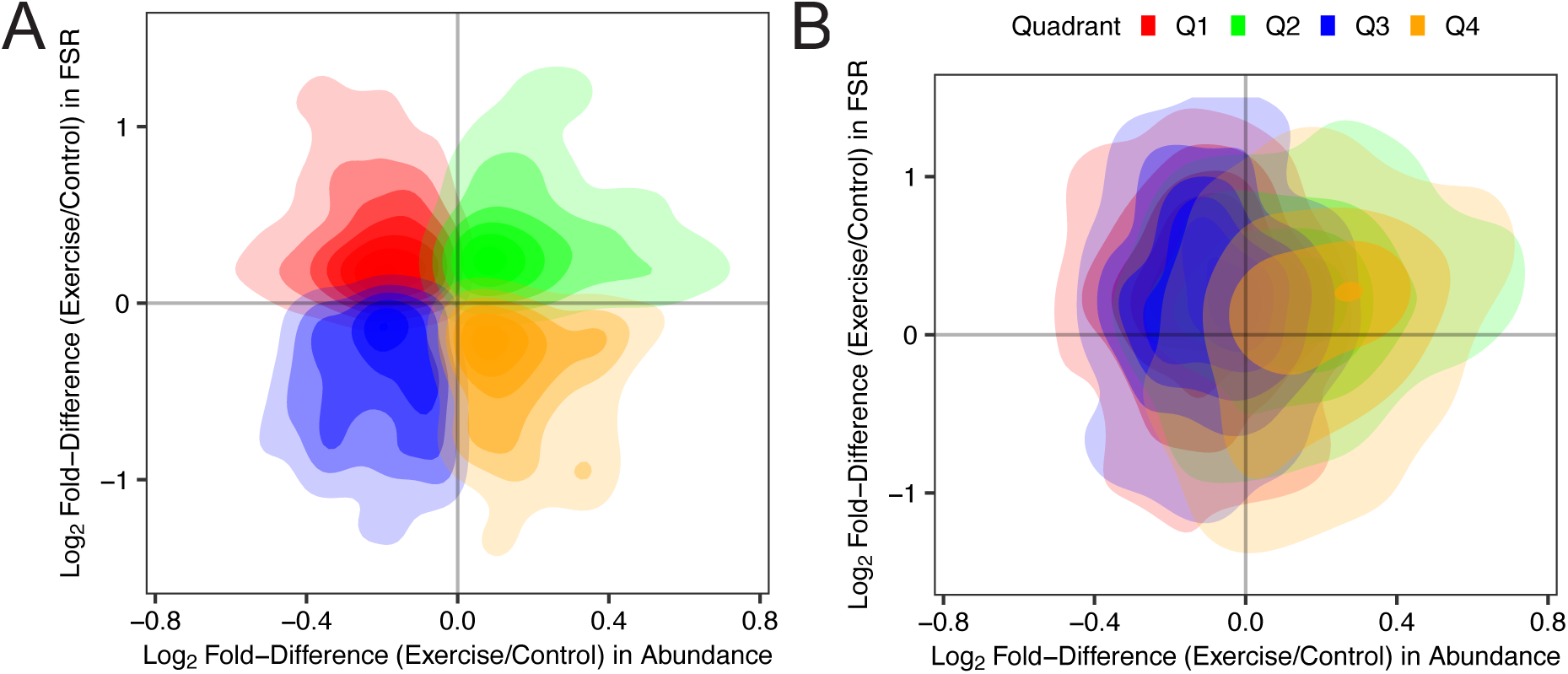
Distinct dynamic proteome responses to resistance exercise between younger and older skeletal muscles (A) Scatter plot visualizing the distribution of proteins based on the log2 fold-difference (Exercise/Control) in abundance (x-axis) and FTR (y-axis) in younger muscle. Data points are categorized into four quadrants based on the direction of fold-difference: Q1 (red) represents proteins with increased FTR but decreased abundance (top-left), Q2 (blue) represents proteins with increased FTR and abundance (top-right), Q3 (green) represents proteins with decreased FTR and abundance (bottom-left), and Q4 (orange) represents proteins with decreased FTR but increased abundance (bottom-right). Density contours are shown to indicate regions of higher protein density within each quadrant. (B) The proteins categorized into four quadrants in younger muscle were mapped based on log2 Fold-Difference (Exercise/Control) in abundance and FTR in older muscle. Horizontal and vertical lines at x = 0 and y = 0 denote no difference in abundance and FTR, respectively.

**Figure 7.**
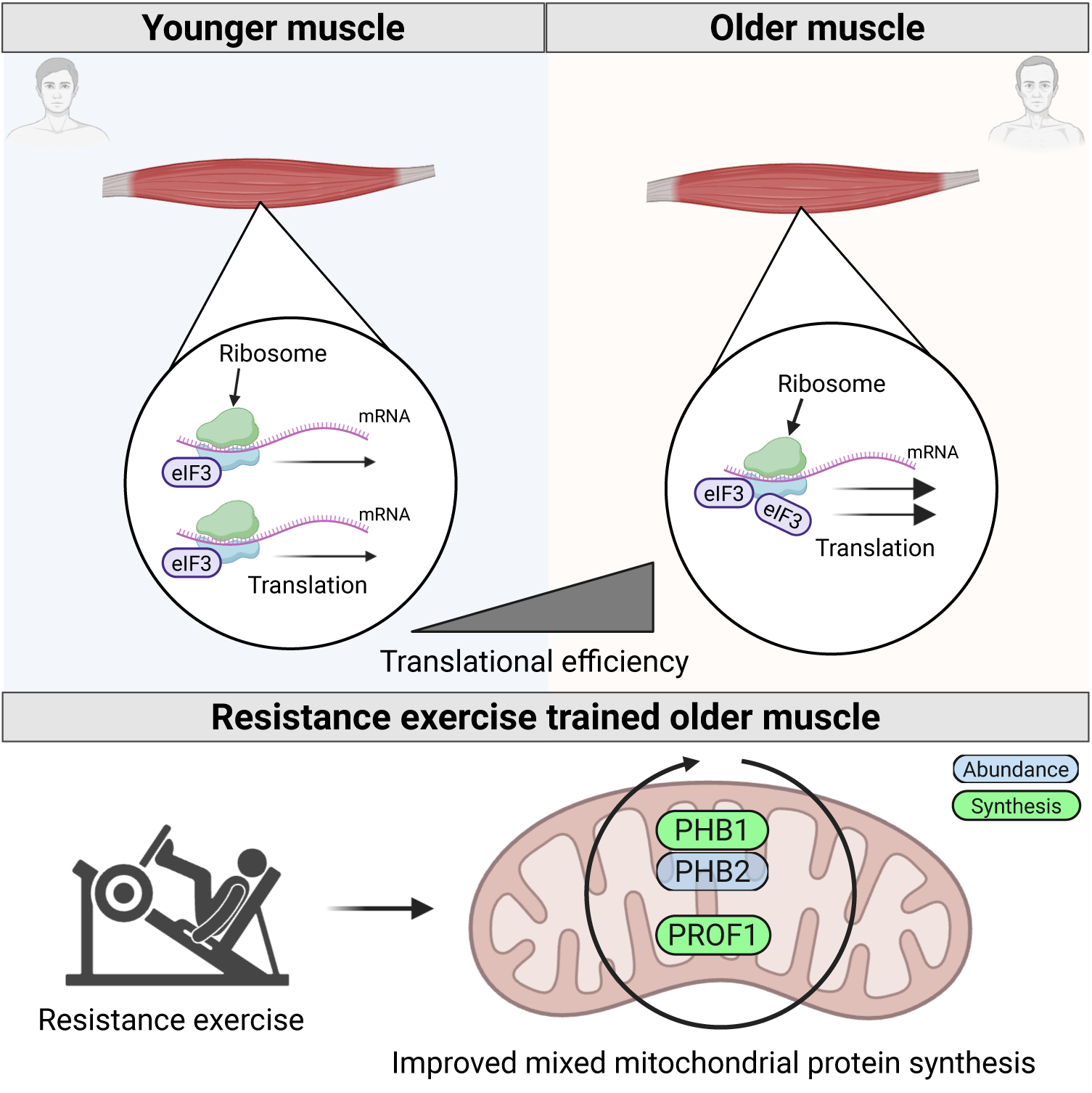
Age-related proteostasis imbalance in skeletal muscle and the restorative effects of resistance exercise *Dynamic Proteome Profiling* revealed a stoichiometric imbalance within the proteostasis network in aged skeletal muscle, including subunits of eIF3, subunits of 40S and 60S ribosomal proteins. However, mixed-muscle protein synthesis was not impaired in older muscle as compared to younger muscle, indicating translational activity might be more efficient in older muscle as compared to younger muscle. Resistance exercise specifically improved mixed mitochondrial protein synthesis in older muscle. Moreover, older muscle specifically increased protein abundance of PHB2 and improved synthesis rate of PHB1 and PROF1, suggesting resistance exercise promotes mitochondrial proteome remodeling. Created in BioRender. Nishimura, Y. (2025) https://BioRender.com/p2a1aio

## Discussion

The preservation of skeletal muscle is a cornerstone of healthy ageing and the use of resistance exercise to enhance muscle protein synthesis in older adults has been a prominent focus of research attention (35). *Dynamic Proteome Profiling* brings new detail to studies on human muscle protein synthesis by simultaneously reporting abundance and FTR data on a protein-by-protein basis. Previous *Dynamic Proteome Profiling* studies in humans uncovered novel protein-specific responses to exercise in younger adults (25) and highlighted losses in muscle proteostasis in the muscle of people with obesity (24), which were hidden from traditional analyses on mixed-protein samples. *Dynamic Proteome Profiling* has not previously been used to investigate the effects of human ageing and, in the current work, *Dynamic Proteome Profiling* discovered ribosomes in the muscle of older men are either more efficient or are ‘harder-worked’ than ribosomes in the muscle of younger adults.

In humans, ageing is associated with a general loss in ribosomal subunits from the muscle proteome (Ubaida-Mohien et al., 2019), and this effect is evident in both fast- and slow-twitch fibers (Murgia et al., 2017). Ubaida-Mohien et al. (Ubaida-Mohien et al., 2019) report significant negative correlations between age (58 health participants aged, 20-87 years) and the abundance of protein subunits of the 40S and 60S ribosome. Similarly, we identified 27 ribosomal proteins (eight 40S proteins and nineteen 60S proteins), which were less abundant in the muscle of older participants. Nevertheless, the rate of bulk, mixed-protein synthesis was not different between younger and older men (Fig. 3 A), which suggests ribosomes in older muscle may exhibit greater translational efficiency to maintain similar levels of protein turnover compared to ribosomes in younger muscle. Protein turnover is necessary for the maintenance of proteome health, but errors in the translation of new protein represent a major burden to the proteostasis network. In an earlier study (Brown et al., 2022), we report replicatively aged C2C12 muscle cells have a lesser abundance of ribosomal proteins and that protein turnover in replicatively aged muscle cells fails to respond appropriately during differentiation compared to non- replicatively aged cells. Therefore, although translational efficiency may be maintained in older muscle through compensatory ribosome activity, there is a risk that translational fidelity may be compromised, potentially resulting in the production of aberrant proteins.

Subunits of the eukaryotic initiation factor 3 complex (eIF3) defied the general pattern of age-related decline exhibited by ribosomal subunits and other eIF complexes, which may indicate alterations in ribosomal heterogeneity. A change in the stoichiometry between ribosomal proteins and subunits of the eIF3 complex may underpin the greater efficiency of older muscle and may favor the translation of mRNA transcripts for mitochondrial proteins. During its canonical function, the eIF3 complex separates from 43S pre-initiation prior to formation of the 80S ribosome, however, eIF3 can also be retained and continue to interact with 80S ribosome to aid the elongation of challenging pre-sequences which are common in the transcripts for mitochondrial proteins (65). Skeletal muscle mitochondrial protein abundance (31), synthesis rate (66), and function (33) decline with ageing, and polysome profiling (67) suggests the muscle of older adults may deprioritize the translation of mitochondrial proteins. In the current analysis, subunits of mitochondrial respiratory complexes were less abundant in the muscle proteome of older adults. Resistance exercise training is capable of increasing mitochondrial protein abundance in the muscle of older adults (33, 68) when conducted over a 12-week period. The 15-day intervention reported here did not change mitochondrial protein abundance but did tend to increase the turnover rate of mitochondrially annotated proteins specifically in older participants (Fig. 4B) and some individual mitochondrial proteins exhibited significant age-specific responses to resistance training. The abundance of PHB2 and FTR of PHB1 and PROF1 were specifically increased in older muscle during resistance exercise. A recent study discovered that PROF1 is localized within mitochondria and maintains mitochondrial integrity (69). PHB1 and PHB2 are proteins of the inner mitochondrial membrane and assemble into large ring complexes that stabilize newly synthesized mitochondrial translation products (70). Moreover, PHB2 is a recognized mitophagy receptor (71) when exposed on the outer membrane of mitochondria and, thereby, further regulates mitochondrial proteostasis. The unique resistance exercise response of mitochondrial proteins in older skeletal muscle may highlight the greater demand of the PHB complex to stabilize those newly synthesized mitochondrial proteins, implicating an improved mitochondrial proteostasis in older muscle.

The initial bouts of a new regimen of resistance training can disrupt sarcomere structure (known as Z-band streaming) and are associated with acute elevations in the rate of mixed-protein turnover in human muscle (72). The Z-disk of striated muscle is amongst the most densely packed protein structures in eukaryotic cells (73) and is a phosphorylation hotspot in myofibrils after contractile work (74). Putative mechanosensory proteins, including SPEGβ (75), CSRP3 (76) and FLNC can localize to the Z-band and were responsive to resistance training (Fig. 4 B). CSRP3 was more abundant in younger adults, and the FTR of CSRP3 increased in response to resistance exercise in both younger and older men. The turnover rate of FLNC also increased in response to resistance training in both older and younger adults. FLNC is an actin binding protein and dimers of FLNC contribute to the 3-dimensional F-actin structure of the Z-disc. In addition, FLNC contains mechanosensitive elastic regions that open under contractile force, to expose immunoglobulin repeat domains that interact with other proteins (77). Razinia et al. (78) reports filamin proteins interact with ASB2β, which is a subunit of a substrate-recognition component of a SCF-like ECS (Elongin-Cullin-SOCS-box protein) ubiquitin E3 ligase complex. In the current work, ASB2β was more abundant in the muscle of older adults at baseline and decreased in abundance specifically in older muscle after resistance exercise training. Overexpression of ASB2β induces muscle atrophy, which is accompanied by gains in the abundance of FLNC and other sarcomere and Z-disk proteins (60).

As expected, older men exhibited whole-body insulin resistance (Table 1), and some age-related differences in the muscle proteome were associated with muscle metabolism and metabolic regulatory pathways. Adiponectin (ADIPO), which is an adipokine associated with whole-body insulin sensitivity (79), and Copine-3 (CPNE3), which regulates glucose-stimulated insulin secretion in pancreatic-β cells (80), were amongst the proteins that increased in FTR in response to resistance training specifically in the muscle of younger adults. Protein feeding and amino acid availability are also important components of muscle metabolism linked to ageing (81, 82). Proteins involved in the regulation of the mTORC1 by amino acids were amongst the proteins highlighted in our analyses. The leucine sensor, SESN1 was more abundant in older muscle at baseline, whereas LTOR1, which tethers the Ragulator complex to the lysosome (83), was decreased in older and increased in younger muscle in response to resistance exercise. When bound with leucine, SESN1 is released from GATOR2 and activates mTORC1 signaling in skeletal muscle (59), and overexpression of SESN1 in older mice protects against age-related muscle atrophy (84).

Our current analysis of 1046 protein-specific FTR values represents the largest collection of proteins studied in human skeletal muscle using D2O-based *Dynamic Proteome Profiling* (23–25, 85, 86). However, this pilot study used a small sample size (n=4 per group) and was conducted in males only. The characteristics of muscle ageing are different between men and women (56, 87) and ageing is also a continuous and likely a non-linear process (88), which cannot be captured by our two-group design. Muscle adaptation to resistance training is also a time-dependent process (27), which cannot be fully captured from comparisons between baseline and only a single post-intervention time point. In the future, longer periods of D2O labelling and timeseries sampling as performed in Camera et al. (25) will be required to add further understanding of age-related differences in the muscle response to resistance training.

## Conclusion

In conclusion, age-related differences in muscle protein abundance profile were associated with different dynamic muscle proteome responses to resistance exercise between older and younger males. Actin binding proteins associated with the Z-Disc were particularly enriched amongst exercise responsive proteins in younger and older men, whereas resistance exercise specifically improved the turnover of mitochondrial proteins in older muscle. Mixed protein turnover was similar between younger and older adults, but the abundance of subunits of the small- and large- ribosomal complexes was less in the muscle of older men. These early data provide an impetus for further detailed exploration of muscle proteostasis in human ageing and the use of resistance exercise and *Dynamic Proteome Profiling* as tools to find new therapeutic strategies to maintain muscle health.

## Data availability

All data needed to evaluate the conclusions or reperform analyses in the paper are present in the paper or the Supplementary Materials. The mass spectrometry proteomics data generated in this study have been deposited to the ProteomeXchange Consortium via the PRIDE (76) under the dataset identifiers PXD062373 and 10.6019/PXD062373.

## Supplemental Data

This article contains supplemental data.

## Funding and additional information

The research was supported by the ART (Ageing Research Translation) of Healthy Ageing Network which is funded by UK Research and Innovation (Grant Ref: BB/W018209/1). YN was supported by the Physiological Society UK via the Unlocking Futures Fund.

## Author contributions (CRediT)

Stewart, Burniston: Conceptualization (Ideas; formulation or evolution of overarching research goals and aims)

Nishimura, Rudolf, Barrett, Kirwan, Johnson, Pugh, Strauss, Areta, Owens, Jackson, Foster, Ortega-Martorell, Khaiyat, Stewart, Akpan, Burniston: Methodology (Development or design of methodology; creation of models)

Nishimura, Stead, Burniston: Software (Programming, software development; designing computer programs; implementation of the computer code and supporting algorithms; testing of existing code components)

Nishimura, Rudolf, Barrett, Kirwan, Johnson, Pugh, Strass, Areta, Stead, Owens: Validation (Verification, whether as a part of the activity or separate, of the overall replication/ reproducibility of results/experiments and other research outputs)

Nishimura, Rudolf, Barrett, Stead, Burniston: Formal analysis (Application of statistical, mathematical, computational, or other formal techniques to analyze or synthesize study data)

Nishimura, Rudolf, Barrett, Kirwan, Johnson, Pugh, Strauss, Areta, Owens, Jackson: Investigation (Conducting a research and investigation process, specifically performing the experiments, or data/evidence collection)

Pugh, Strauss, Areta, Owens, Jackson, Foster, Khaiyat, Burniston: Resources (Provision of study materials, reagents, materials, patients, laboratory samples, animals, instrumentation, computing resources, or other analysis tools)

Nishimura, Rudolf, Barrett, Kirwan, Johnson, Jackson, Foster, Burniston: Data Curation (Management activities to annotate (produce metadata), scrub data and maintain research data (including software code, where it is necessary for interpreting the data itself) for initial use and later reuse)

Nishimura, Rudolf, Barrett, Stead, Burniston: Writing - Original Draft (Preparation, creation and/or presentation of the published work, specifically writing the initial draft)

Nishimura, Rudolf, Barrett, Kirwan, Johnson, Pugh, Strauss, Areta, Owens, Jackson, Foster, Ortega-Martorell, Khaiyat, Stewart, Akpan, Burniston: Writing - Review & Editing (Preparation, creation and/or presentation of the published work by those from the original research group, specifically critical review, commentary or revision – including pre-or post-publication stages)

Nishimura, Rudolf, Barrett, Burniston: Visualization (Preparation, creation and/or presentation of the published work, specifically visualization/ data presentation)

Ortega-Martorell, Khaiyat, Stewart, Akpan, Burniston: Supervision (Oversight and leadership responsibility for the research activity planning and execution, including mentorship external to the core team)

Foster, Ortega-Martorell, Khaiyat, Stewart, Akpan, Burniston: Project administration (Management and coordination responsibility for the research activity planning and execution)

Nishimura, Rudolf, Barrett, Kirwan, Johnson, Pugh, Strauss, Areta, Owens, Jackson, Foster, Ortega-Martorell, Khaiyat, Stewart, Akpan, Burniston: Funding acquisition (Acquisition of the financial support for the project leading to this publication)

## Conflict of interest

The authors declare that they have no known competing financial interests or personal relationships that could have appeared to influence the work reported in this paper.

## Notes

### Competing Interest Statement

The authors have declared no competing interest.

## References

1. Christian, C. J., and Benian, G. M. (2020) Animal models of sarcopenia. Aging cell 19, e13223

2. Metter, E. J., Talbot, L., A, Schrager, M., and Conwit, R. (2002) Skeletal muscle strength as a predictor of all-cause mortality in healthy men. J Gerontol A Biol Sci Med Sci 57A, B359-B365

3. Ruiz, J. R., Sui, X., Lobelo, F., Morrow, J. R., Jr., Jackson, A. W., Sjostrom, M., and Blair, S. N. (2008) Association between muscular strength and mortality in men: prospective cohort study. BMJ 337, a439

4. Veronese, N., Koyanagi, A., Cereda, E., Maggi, S., Barbagallo, M., Dominguez, L. J., and Smith, L. (2022) Sarcopenia reduces quality of life in the long-term: longitudinal analyses from the English longitudinal study of ageing. Eur Geriatr Med 13, 633–639

5. Cruz-Jentoft, A. J., Bahat, G., Bauer, J., Boirie, Y., Bruyere, O., Cederholm, T., Cooper, C., Landi, F., Rolland, Y., Sayer, A. A., Schneider, S. M., Sieber, C. C., Topinkova, E., Vandewoude, M., Visser, M., Zamboni, M., Writing Group for the European Working Group on Sarcopenia in Older, P., and the Extended Group for, E. (2019) Sarcopenia: revised European consensus on definition and diagnosis. Age Ageing 48, 16-31

6. Lopez-Otin, C., Blasco, M. A., Partridge, L., Serrano, M., and Kroemer, G. (2023) Hallmarks of aging: An expanding universe. Cell 186, 243–278

7. Kaushik, S., and Cuervo, A. M. (2015) Proteostasis and aging. Nature Medicine 21, 1406–1415

8. Wiśniewski, J. R., Hein, M. Y., Cox, J., and Mann, M. (2014) A "proteomic ruler" for protein copy number and concentration estimation without spike-in standards. Molecular and Cellular Proteomics 13, 3497–3506

9. Hohfeld, J., Benzing, T., Bloch, W., Furst, D. O., Gehlert, S., Hesse, M., Hoffmann, B., Hoppe, T., Huesgen, P. F., Kohn, M., Kolanus, W., Merkel, R., Niessen, C. M., Pokrzywa, W., Rinschen, M. M., Wachten, D., and Warscheid, B. (2021) Maintaining proteostasis under mechanical stress. EMBO Rep 22, e52507

10. Lopez-Otin, C., Blasco, M. A., Partridge, L., Serrano, M., and Kroemer, G. (2013) The hallmarks of aging. Cell 153, 1194–1217

11. Balch, W. E., Morimoto, R. I., Dillin, A., and Kelly, J. W. (2008) Adapting proteostasis for disease intervention. Science 319, 916–919

12. Demontis, F., Piccirillo, R., Goldberg, A. L., and Perrimon, N. (2013) Mechanisms of skeletal muscle aging: insights from Drosophila and mammalian models. Dis Model Mech 6, 1339–1352

13. Granic, A., Suetterlin, K., Shavlakadze, T., Grounds, M. D., and Sayer, A. A. (2023) Hallmarks of ageing in human skeletal muscle and implications for understanding the pathophysiology of sarcopenia in women and men. Clin Sci (Lond) 137, 1721–1751

14. Herndon, L. A., Schmeissner, P. J., Dudaronek, J. M., Brown, P. A., Listner, K. M., Sakano, Y., Paupard, M. C., Hall, D. H., and Driscoll, M. (2002) Stochastic and genetic factors influence tissue-specific decline in ageing C. elegans. Nature 419, 808–814

15. Walther, D. M., Kasturi, P., Zheng, M., Pinkert, S., Vecchi, G., Ciryam, P., Morimoto, R. I., Dobson, C. M., Vendruscolo, M., Mann, M., and Hartl, F. U. (2015) Widespread proteome remodeling and aggregation in aging C. elegans. Cell 161, 919–932

16. Dhondt, I., Petyuk, V. A., Bauer, S., Brewer, H. M., Smith, R. D., Depuydt, G., and Braeckman, B. P. (2017) Changes of protein turnover in aging caenorhabditis elegans. Molecular and Cellular Proteomics 16, 1621–1633

17. Volpi, E., Sheffield-Moore, M., Rasmussen, B. B., and Wolfe, R. R. (2001) Basal muscle amino acid kinetics and protein synthesis in healthy young and older men. Jama 286, 1206–1212

18. Brook, M. S., Wilkinson, D. J., Mitchell, W. K., Lund, J. N., Phillips, B. E., Szewczyk, N. J., Greenhaff, P. L., Smith, K., and Atherton, P. J. (2016) Synchronous deficits in cumulative muscle protein synthesis and ribosomal biogenesis underlie age-related anabolic resistance to exercise in humans. J Physiol 594, 7399–7417

19. Marshall, R. N., Morgan, P. T., Smeuninx, B., Quinlan, J. I., Brook, M. S., Atherton, P. J., Smith, K., Wilkinson, D. J., and Breen, L. (2023) Myofibrillar Protein Synthesis and Acute Intracellular Signaling with Elastic Band Resistance Exercise in Young and Older Men. Med Sci Sports Exerc 55, 398–408

20. Cuthbertson, D. (2004) Anabolic signaling deficits underlie amino acid resistance of wasting, aging muscle. The FASEB Journal, 422-424

21. Kumar, V., Selby, A., Rankin, D., Patel, R., Atherton, P., Hildebrandt, W., Williams, J., Smith, K., Seynnes, O., Hiscock, N., and Rennie, M. J. (2009) Age-related differences in the dose-response relationship of muscle protein synthesis to resistance exercise in young and old men. J Physiol 587, 211–217

22. Short, K. R., Vittone, J. L., Bigelow, M. L., Proctor, D. N., and Nair, K. S. (2004) Age and aerobic exercise training effects on whole body and muscle protein metabolism. American journal of physiology. Endocrinology and metabolism 286, E92–101

23. Stansfield, B. N., Barrett, J. S., Bennett, S., Stead, C. A., Pugh, J., Shepherd, S. O., Strauss, J. A., Louis, J., Close, G. L., and Lisboa, P. J. (2025) Turnover rates of human muscle proteins in vivo reported in fractional, mole and absolute units. BMC Methods 2, 9

24. Srisawat, K., Stead, C. A., Hesketh, K., Pogson, M., Strauss, J. A., Cocks, M., Siekmann, I., Phillips, S. M., Lisboa, P. J., Shepherd, S., and Burniston, J. G. (2023) People with obesity exhibit losses in muscle proteostasis that are partly improved by exercise training. *Proteomics*, e2300395

25. Camera, D. M., Burniston, J. G., Pogson, M. A., Smiles, W. J., and Hawley, J. A. (2017) Dynamic proteome profiling of individual proteins in human skeletal muscle after a high-fat diet and resistance exercise. FASEB J 31, 5478–5494

26. Hesketh, S. J., Sutherland, H., Lisboa, P. J., Jarvis, J. C., and Burniston, J. G. (2020) Adaptation of rat fast-twitch muscle to endurance activity is underpinned by changes to protein degradation as well as protein synthesis. FASEB J 34, 10398–10417

27. Stead, C. A., Hesketh, S. J., Thomas, A. C. Q., Viggars, M. R., Sutherland, H., Jarvis, J. C., and Burniston, J. G. (2025) Dynamic time course of muscle proteome adaptation to programmed resistance training in rats. Am J Physiol Cell Physiol

28. Murgia, M., Toniolo, L., Nagaraj, N., Ciciliot, S., Vindigni, V., Schiaffino, S., Reggiani, C., and Mann, M. (2017) Single Muscle Fiber Proteomics Reveals Fiber-Type-Specific Features of Human Muscle Aging. Cell Reports 19, 2396–2409

29. Deane, C. S., Phillips, B. E., Willis, C. R. G., Wilkinson, D. J., Smith, K., Higashitani, N., Williams, J. P., Szewczyk, N. J., Atherton, P. J., Higashitani, A., and Etheridge, T. (2022) Proteomic features of skeletal muscle adaptation to resistance exercise training as a function of age. *GeroScience*

30. Theron, L., Gueugneau, M., Coudy, C., Viala, D., Bijlsma, A., Butler-Browne, G., Maier, A., Bechet, D., and Chambon, C. (2014) Label-free quantitative protein profiling of vastus lateralis muscle during human aging. Mol Cell Proteomics 13, 283–294

31. Ubaida-Mohien, C., Lyashkov, A., Gonzalez-Freire, M., Tharakan, R., Shardell, M., Moaddel, R., Semba, R. D., Chia, C. W., Gorospe, M., Sen, R., and Ferrucci, L. (2019) Discovery proteomics in aging human skeletal muscle finds change in spliceosome, immunity, proteostasis and mitochondria. eLife 8, 1–27

32. Adelnia, F., Ubaida-Mohien, C., Moaddel, R., Shardell, M., Lyashkov, A., Fishbein, K. W., Aon, M. A., Spencer, R. G., and Ferrucci, L. (2020) Proteomic signatures of in vivo muscle oxidative capacity in healthy adults. Aging Cell 19, 1–12

33. Robinson, M. M., Dasari, S., Konopka, A. R., Johnson, M. L., Manjunatha, S., Esponda, R. R., Carter, R. E., Lanza, I. R., and Nair, K. S. (2017) Enhanced Protein Translation Underlies Improved Metabolic and Physical Adaptations to Different Exercise Training Modes in Young and Old Humans. Cell Metabolism 25, 581–592

34. Stead, C. A., Thomas, A., Nishimura, Y., Abbasi, M., Barrett, J., and Burniston, J. G. (2025) Muscle Proteome Dynamics. *The Skeletal Muscle: Plasticity*, Degeneration and Epigenetics, 113-153

35. McKendry, J., Stokes, T., McLeod, J. C., and Phillips, S. M. (2021) Resistance Exercise, Aging, Disuse, and Muscle Protein Metabolism. Compr Physiol 11, 2249–2278

36. Nishimura, Y., Bittel, A. J., Stead, C. A., Chen, Y.-W., and Burniston, J. G. (2023) Facioscapulohumeral muscular dystrophy is associated with altered myoblast proteome dynamics. Molecular & Cellular Proteomics

37. Bohannon, R. W. (2006) Reference values for the five-repetition sit-to-stand test: a descriptive meta-analysis of data from elders. Perceptual and motor skills 103, 215–222

38. Balogun, J., and Onigbinde, A. (1991) Intratester reliability and validity of the takei kiki kogo hand grip dynamometer. J Phys Ther Sci 3, 55–60

39. Yu, R., Ong, S., Cheung, O., Leung, J., and Woo, J. (2017) Reference Values of Grip Strength, Prevalence of Low Grip Strength, and Factors Affecting Grip Strength Values in Chinese Adults. J Am Med Dir Assoc 18, 551.e559–551.e516

40. Habets, B., Staal, J. B., Tijssen, M., and van Cingel, R. (2018) Intrarater reliability of the Humac NORM isokinetic dynamometer for strength measurements of the knee and shoulder muscles. BMC Research Notes 11, 15

41. Srisawat, K., Hesketh, K., Cocks, M., Strauss, J., Edwards, B. J., Lisboa, P. J., Shepherd, S., and Burniston, J. G. (2020) Reliability of protein abundance and synthesis measurements in human skeletal muscle. Proteomics 20, 1900194

42. Matsuda, M., and DeFronzo, R. A. (1999) Insulin sensitivity indices obtained from oral glucose tolerance testing: Comparison with the euglycemic insulin clamp. Diabetes Care 22, 1462–1470

43. Wiśniewski, J. R., Zougman, A., Nagaraj, N., and Mann, M. (2009) Universal sample preparation method for proteome analysis. Nature Methods 6, 359–362

44. Nishimura, Y., Bittel, A., Jagan, A., Chen, Y.-W., and Burniston, J. (2025) Proteomic profiling uncovers sexual dimorphism in the muscle response to wheel running exercise in the FLExDUX4 murine model of facioscapulohumeral muscular dystrophy. Molecular & Cellular Proteomics, 101013

45. Silva, J. C., Gorenstein, M. V., Li, G.-z., Vissers, J. P. C., and Geromanos, S. J. (2006) Absolute Quantification of Proteins by LCMS E. Molecular & Cellular Proteomics 5, 144–156

46. Sadygov, R. G. (2020) Partial Isotope Profiles Are Sufficient for Protein Turnover Analysis Using Closed-Form Equations of Mass Isotopomer Dynamics. Anal Chem 92, 14747–14753

47. Holmes, W. E., Angel, T. E., Li, K. W., and Hellerstein, M. K. (2015) Dynamic Proteomics: In Vivo Proteome-Wide Measurement of Protein Kinetics Using Metabolic Labeling. pp. 219–276

48. Ilchenko, S., Haddad, A., Sadana, P., Recchia, F. A., Sadygov, R. G., and Kasumov, T. (2019) Calculation of the Protein Turnover Rate Using the Number of Incorporated 2H Atoms and Proteomics Analysis of a Single Labeled Sample. Analytical Chemistry 91, 14340–14351

49. Bates, D., Mächler, M., Bolker, B., and Walker, S. (2015) Fitting Linear Mixed-Effects Models Usinglme4. Journal of Statistical Software 67

50. Storey, J. D., and Tibshirani, R. (2003) Statistical significance for genomewide studies. Proceedings of the National Academy of Sciences of the United States of America 100, 9440–9445

51. Zhang, B., Kirov, S., and Snoddy, J. (2005) WebGestalt: an integrated system for exploring gene sets in various biological contexts. Nucleic Acids Res 33, W741–748

52. Eden, E., Navon, R., Steinfeld, I., Lipson, D., and Yakhini, Z. (2009) GOrilla: a tool for discovery and visualization of enriched GO terms in ranked gene lists. BMC bioinformatics 10, 1–7

53. Rath, S., Sharma, R., Gupta, R., Ast, T., Chan, C., Durham, T. J., Goodman, R. P., Grabarek, Z., Haas, M. E., Hung, W. H. W., Joshi, P. R., Jourdain, A. A., Kim, S. H., Kotrys, A. V., Lam, S. S., McCoy, J. G., Meisel, J. D., Miranda, M., Panda, A., Patgiri, A., Rogers, R., Sadre, S., Shah, H., Skinner, O. S., To, T. L., Walker, M. A., Wang, H., Ward, P. S., Wengrod, J., Yuan, C. C., Calvo, S. E., and Mootha, V. K. (2021) MitoCarta3.0: an updated mitochondrial proteome now with sub-organelle localization and pathway annotations. Nucleic Acids Res 49, D1541–D1547

54. Pham, T. V., Henneman, A. A., and Jimenez, C. R. (2020) Iq: An R package to estimate relative protein abundances from ion quantification in DIA-MS-based proteomics. Bioinformatics 36, 2611–2613

55. Gayoso-Diz, P., Otero-Gonzalez, A., Rodriguez-Alvarez, M. X., Gude, F., Cadarso-Suarez, C., García, F., and De Francisco, A. (2011) Insulin resistance index (HOMA-IR) levels in a general adult population: curves percentile by gender and age. The EPIRCE study. Diabetes Res Clin Pract 94, 146–155

56. Metter, E. J., Lynch, N., Conwit, R., Lindle, R., Tobin, J., and Hurley, B. (1999) Muscle quality and age: Cross-sectional and longitudinal comparisons. Journals of Gerontology - Series A Biological Sciences and Medical Sciences 54, 8207–8218

57. Wrucke, D. J., Kuplic, A., Adam, M. D., Hunter, S. K., and Sundberg, C. W. (2024) Neural and muscular contributions to the age-related differences in peak power of the knee extensors in men and women. Journal of Applied Physiology 137, 1021–1040

58. Osawa, Y., Studenski, S. A., and Ferrucci, L. (2018) Knee extension rate of velocity development affects walking performance differently in men and women. Experimental Gerontology 112, 63–67

59. Xu, D., Shimkus, K. L., Lacko, H. A., Kutzler, L., Jefferson, L. S., and Kimball, S. R. (2019) Evidence for a role for Sestrin1 in mediating leucine-induced activation of mTORC1 in skeletal muscle. Am J Physiol Endocrinol Metab 316, E817–e828

60. Goodman, C. A., Davey, J. R., Hagg, A., Parker, B. L., and Gregorevic, P. (2021) Dynamic Changes to the Skeletal Muscle Proteome and Ubiquitinome Induced by the E3 Ligase, ASB2beta. *Mol Cell Proteomics* 20, 100050

61. Sharma, A., Zehra, A., and Mathew, S. J. (2024) Myosin heavy chain-perinatal regulates skeletal muscle differentiation, oxidative phenotype and regeneration. The FEBS Journal 291, 2836–2848

62. Vainberg, I. E., Lewis, S. A., Rommelaere, H., Ampe, C., Vandekerckhove, J., Klein, H. L., and Cowan, N. J. (1998) Prefoldin, a chaperone that delivers unfolded proteins to cytosolic chaperonin. Cell 93, 863–873

63. Wu, S. A., Li, Z. J., and Qi, L. (2025) Endoplasmic reticulum (ER) protein degradation by ER-associated degradation and ER-phagy. Trends in Cell Biology

64. Sylow, L., Jensen, T. E., Kleinert, M., Mouatt, J. R., Maarbjerg, S. J., Jeppesen, J., Prats, C., Chiu, T. T., Boguslavsky, S., Klip, A., Schjerling, P., and Richter, E. A. (2013) Rac1 is a novel regulator of contraction-stimulated glucose uptake in skeletal muscle. Diabetes 62, 1139–1151

65. Lin, Y., Li, F., Huang, L., Polte, C., Duan, H., Fang, J., Sun, L., Xing, X., Tian, G., Cheng, Y., Ignatova, Z., Yang, X., and Wolf, D. A. (2020) eIF3 Associates with 80S Ribosomes to Promote Translation Elongation, Mitochondrial Homeostasis, and Muscle Health. Mol Cell 79, 575–587 e577

66. Rooyackers, O. E., Adey, D. B., Ades, P. A., and Nair, K. S. (1996) Effect of age on in vivo rates of mitochondrial protein synthesis in human skeletal muscle. Proceedings of the National Academy of Sciences of the United States of America 93, 15364–15369

67. Tharakan, R., Ubaida-Mohien, C., Piao, Y., Gorospe, M., and Ferrucci, L. (2021) Ribosome profiling analysis of human skeletal muscle identifies reduced translation of mitochondrial proteins with age. RNA Biol 18, 1555–1559

68. Mesquita, P. H. C., Lamb, D. A., Parry, H. A., Moore, J. H., Smith, M. A., Vann, C. G., Osburn, S. C., Fox, C. D., Ruple, B. A., Huggins, K. W., Fruge, A. D., Young, K. C., Kavazis, A. N., and Roberts, M. D. (2020) Acute and chronic effects of resistance training on skeletal muscle markers of mitochondrial remodeling in older adults. Physiol Rep 8, e14526

69. Sun, Y., Pang, Y., Wu, X., Zhu, R., Wang, L., Tian, M., He, X., Liu, D., and Yang, X. (2024) Landscape of alternative splicing and polyadenylation during growth and development of muscles in pigs. Commun Biol 7, 1607

70. Nijtmans, L. G., de Jong, L., Sanz, M. A., Coates, P. J., Berden, J. A., Back, J. W., Muijsers, A. O., van der Spek, H., and Grivell, L. A. (2000) Prohibitins act as a membrane-bound chaperone for the stabilization of mitochondrial proteins. The EMBO journal 19, 2444–2451

71. Wei, Y., Chiang, W. C., Sumpter, R., Jr., Mishra, P., and Levine, B. (2017) Prohibitin 2 Is an Inner Mitochondrial Membrane Mitophagy Receptor. Cell 168, 224–238 e210

72. Damas, F., Phillips, S. M., Libardi, C. A., Vechin, F. C., Lixandrão, M. E., Jannig, P. R., Costa, L. A. R., Bacurau, A. V., Snijders, T., Parise, G., Tricoli, V., Roschel, H., and Ugrinowitsch, C. (2016) Resistance training-induced changes in integrated myofibrillar protein synthesis are related to hypertrophy only after attenuation of muscle damage. Journal of Physiology 594, 5209–5222

73. Zou, P., Pinotsis, N., Lange, S., Song, Y.-H., Popov, A., Mavridis, I., Mayans, O. M., Gautel, M., and Wilmanns, M. (2006) Palindromic assembly of the giant muscle protein titin in the sarcomeric Z-disk. Nature 439, 229–233

74. Reimann, L., Wiese, H., Leber, Y., Schwäble, A. N., Fricke, A. L., Rohland, A., Knapp, B., Peikert, C. D., Drepper, F., van der Ven, P. F. M., Radziwill, G., Fürst, D. O., and Warscheid, B. (2017) Myofibrillar Z-discs Are a Protein Phosphorylation Hot Spot with Protein Kinase C (PKCα) Modulating Protein Dynamics. Molecular & Cellular Proteomics 16, 346–367

75. Kusić, D., Connolly, J., Kainulainen, H., Semenova, E. A., Borisov, O. V., Larin, A. K., Popov, D. V., Generozov, E. V., Ahmetov, I. I., Britton, S. L., Koch, L. G., and Burniston, J. G. (2020) Striated muscle-specific serine/threonine-protein kinase beta segregates with high versus low responsiveness to endurance exercise training. Physiological Genomics 52, 35–46

76. Kostek, M. C., Chen, Y.-w., Cuthbertson, D. J., Shi, R., Fedele, M. J., Esser, K. A., and Rennie, M. J. (2007) Gene expression responses over 24 h to lengthening and shortening contractions in human muscle: major changes in CSRP3, MUSTN1, SIX1, and FBXO32. Physiological Genomics 31, 42-52

77. Reimann, L., Schwäble, A. N., Fricke, A. L., Mühlhäuser, W. W., Leber, Y., Lohanadan, K., Puchinger, M. G., Schäuble, S., Faessler, E., and Wiese, H. (2020) Phosphoproteomics identifies dual-site phosphorylation in an extended basophilic motif regulating FILIP1- mediated degradation of filamin-C. Communications biology 3, 253

78. Razinia, Z., Baldassarre, M., Bouaouina, M., Lamsoul, I., Lutz, P. G., and Calderwood, D. A. (2011) The E3 ubiquitin ligase specificity subunit ASB2α targets filamins for proteasomal degradation by interacting with the filamin actin-binding domain. Journal of Cell Science 124, 2631–2641

79. Kubota, N., Terauchi, Y., Yamauchi, T., Kubota, T., Moroi, M., Matsui, J., Eto, K., Yamashita, T., Kamon, J., Satoh, H., Yano, W., Froguel, P., Nagai, R., Kimura, S., Kadowaki, T., and Noda, T. (2002) Disruption of adiponectin causes insulin resistance and neointimal formation. J Biol Chem 277, 25863–25866

80. El-Huneidi, W., Anjum, S., Mohammed, A. K., Unnikannan, H., Saeed, R., Bajbouj, K., Abu-Gharbieh, E., and Taneera, J. (2021) Copine 3 “CPNE3” is a novel regulator for insulin secretion and glucose uptake in pancreatic β-cells. Scientific Reports 11, 20692

81. Nishimura, Y., Musa, I., Holm, L., and Lai, Y.-C. (2021) Recent advances in measuring and understanding the regulation of exercise-mediated protein degradation in skeletal muscle. American journal of physiology. Cell physiology, 6-6

82. Nishimura, Y., Højfeldt, G., Breen, L., Tetens, I., and Holm, L. (2021) Dietary protein requirements and recommendations for healthy older adults: a critical narrative review of the scientific evidence. Nutrition research reviews, 1-17

83. Sancak, Y., Bar-Peled, L., Zoncu, R., Markhard, A. L., Nada, S., and Sabatini, D. M. (2010) Ragulator-rag complex targets mTORC1 to the lysosomal surface and is necessary for its activation by amino acids. Cell 141, 290–303

84. A. Segalés, J., Perdiguero, E., Serrano, A. L., Sousa-Victor, P., Ortet, L., Jardí, M., Budanov, V., Garcia-Prat, L., Sandri, M., and Thomson, D. M. (2020) Sestrin prevents atrophy of disused and aging muscles by integrating anabolic and catabolic signals. Nature communications 11, 189

84. Murphy, C. H., Shankaran, M., Churchward-Venne, T. A., Mitchell, C. J., Kolar, N. M., Burke, L. M., Hawley, J. A., Kassis, A., Karagounis, L. G., Li, K., King, C., Hellerstein, M., and Phillips, S. M. (2018) Effect of resistance training and protein intake pattern on myofibrillar protein synthesis and proteome kinetics in older men in energy restriction. J Physiol 596, 2091–2120

85. Shankaran, M., King, C. L., Angel, T. E., Holmes, W. E., Li, K. W., Colangelo, M., Price, J. C., Turner, S. M., Bell, C., Hamilton, K. L., Miller, B. F., and Hellerstein, M. K. (2016) Circulating protein synthesis rates reveal skeletal muscle proteome dynamics. Journal of Clinical Investigation 126, 288–302

86. Roberts, B. M., Lavin, K. M., Many, G. M., Thalacker-Mercer, A., Merritt, E. K., Bickel, C. S., Mayhew, D. L., Tuggle, S. C., Cross, J. M., Kosek, D. J., Petrella, J. K., Brown, C. J., Hunter, G. R., Windham, S. T., Allman, R. M., and Bamman, M. M. (2018) Human neuromuscular aging: Sex differences revealed at the myocellular level. Exp Gerontol 106, 116–124

87. Shen, X., Wang, C., Zhou, X., Zhou, W., Hornburg, D., Wu, S., and Snyder, M. P. (2024) Nonlinear dynamics of multi-omics profiles during human aging. Nature aging, 1-16

