## Supplemental Fig. for "Dynamic proteome profiling uncovers age-related impairments in proteostasis and the protective effects of resistance exercise in human skeletal muscle"

Ageing; biosynthetic labelling; deuterium oxide; eIF3; ribosomes; heavy water; mitochondria; muscle protein synthesis; proteome dynamics; sarcopenia

### Supplemental Figures

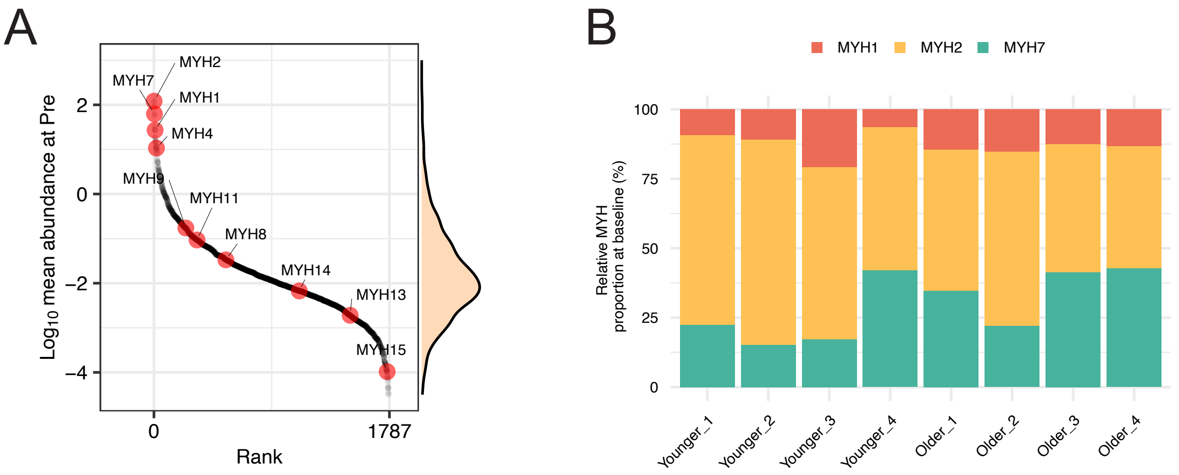


**Figure S1: Myosin heavy chain isoforms are highly abundant in human skeletal muscle with no differences in their proportion between age groups – Related to Fig.2**

(**A**) Rank distribution plot of average protein abundance (n = 1787 proteins) at Pre (day -1). Red data points highlight myosin heavy chain isoforms. (**B**) Bar chart illustrating relative proportion of myosin heavy chain isoforms (MYH1, MYH2, and MYH7) in each participant.

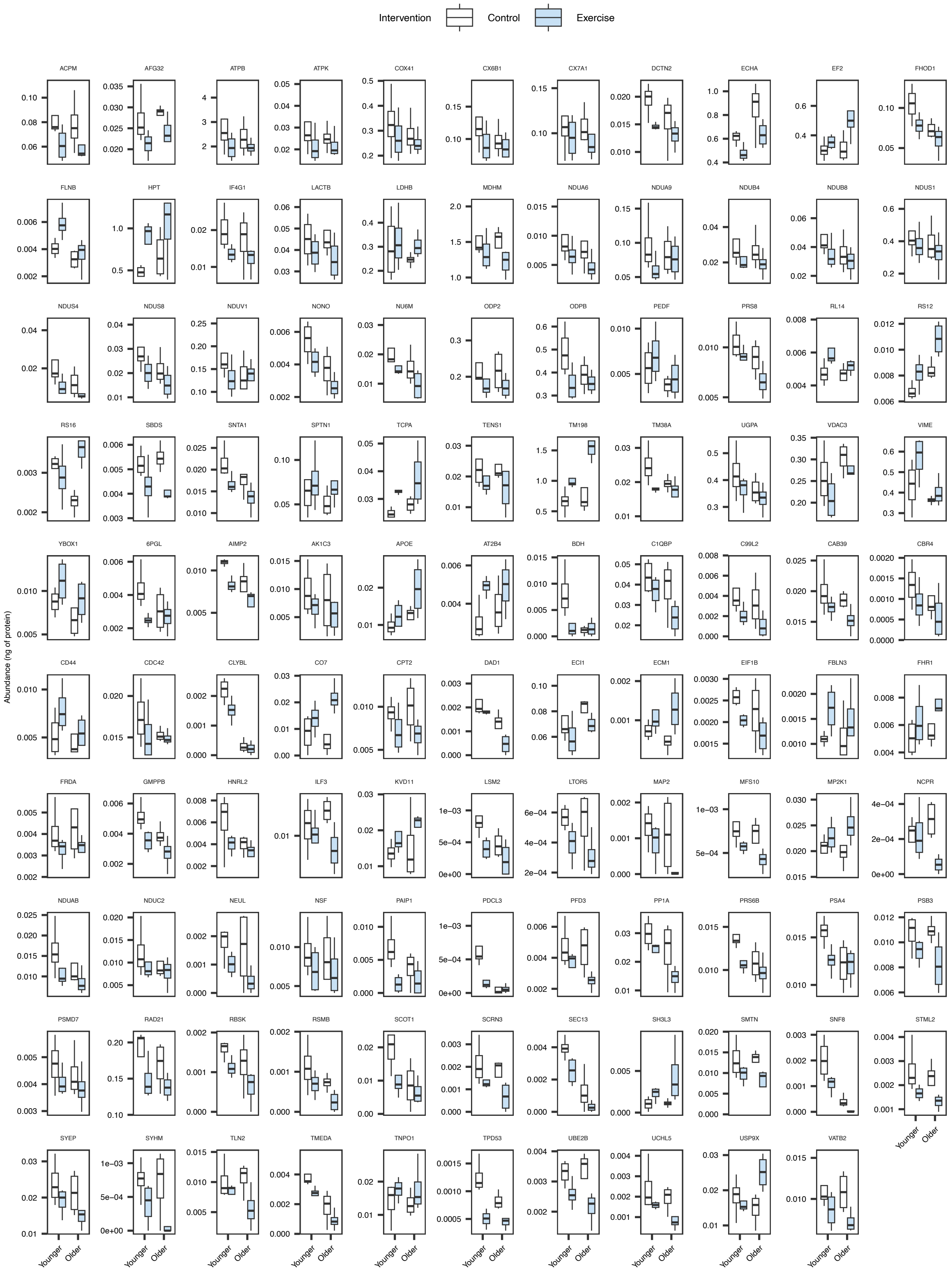


**Figure S2: Box plots of 109 proteins with a main effect of Intervention – Related to Fig.5**

Box plots illustrating differences of protein abundance between Age (Younger versus Older) and Intervention (Control versus Exercise). All $P<0.05$.
